# Unusually Rapid Isomerization of Aspartic Acid in Tau

**DOI:** 10.1101/2024.12.04.626870

**Authors:** Thomas A. Shoff, Brielle Van Orman, Vivian C. Onwudiwe, Joseph C. Genereux, Ryan R. Julian

## Abstract

Spontaneous chemical modifications in long-lived proteins can potentially change protein structure in ways that impact proteostasis and cellular health. For example, isomerization of aspartic acid interferes with protein turnover and is anticorrelated with cognitive acuity in Alzheimer’s disease. However, few isomerization rates have been determined for Asp residues in intact proteins. To remedy this deficiency, we used protein extracts from SH-SY5Y neuroblastoma cells as a source of a complex, brain-relevant proteome with no baseline isomerization. Cell lysates were aged *in vitro* to generate isomers, and extracted proteins were analyzed by data-independent acquisition (DIA) liquid chromatography-mass spectrometry (LC-MS). Although no Asp isomers were detected at Day 0, isomerization increased across time and was quantifiable for 105 proteins by Day 50. Data analysis revealed that isomerization rate is influenced by both primary sequence and secondary structure, suggesting that steric hindrance and backbone rigidity modulate isomerization. Additionally, we examined lysates extracted under gentle conditions to preserve protein complexes and found that protein-protein interactions often slow isomerization. Base catalysis was explored as a means to accelerate Asp isomerization due to findings of accelerated asparagine deamidation. However, no substantial rate enhancement was found for isomerization, suggesting fundamental differences in acid-base chemistry. With an enhanced understanding of Asp isomerization in proteins in general, we next sought to better understand Asp isomerization in tau. *In vitro* aging of monomeric and aggregated recombinant tau revealed that tau isomerizes significantly faster than any similar protein within our dataset, which is likely related to its correlation with cognition in Alzheimer’s disease.

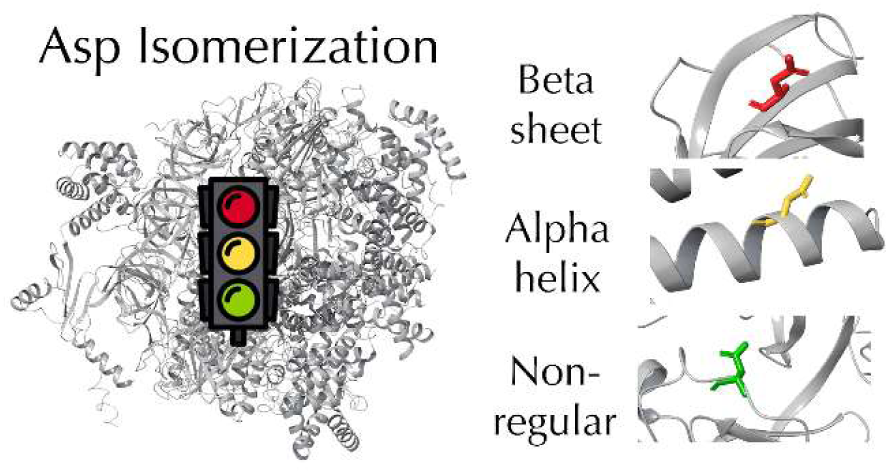

## Introduction

The vast majority of proteins in the human body are synthesized, folded, and degraded back into amino acids within 24 hours to 10 days.^1–3^ However, long-lived proteins (LLPs) are a subset of the proteome that have half-lives ranging from weeks to decades.^4^ LLPs vary in both localization and function, with examples including structural proteins found in neurons, all proteins within the eye lens, and nuclear pore complex proteins, among others.^5–10^ Due to their lengthened presence in cells, LLPs can accumulate extensive post-translational modifications (PTMs) and spontaneous chemical modifications (SCMs). SCMs, including oxidation, isomerization, deamidation, and epimerization, occur in the absence of enzymatic activity,^11–13^ and often lead to changes in protein function, structure, recognition, and turnover.^14–18^

Isomerization of aspartic acid (Asp) is an example of an SCM that has been shown to increase throughout human aging.^8,9^ As shown in Scheme 1, L-Asp can undergo nucleophilic attack of the sidechain by the backbone nitrogen to form a succinimide ring intermediate. This short-lived species opens by hydrolysis to form L-Asp or L-isoAsp. L-succinimide can also stereoisomerize to form both D-Asp and D-isoAsp, leading to a total of four potential isomeric forms.^19^

**Scheme 1.**
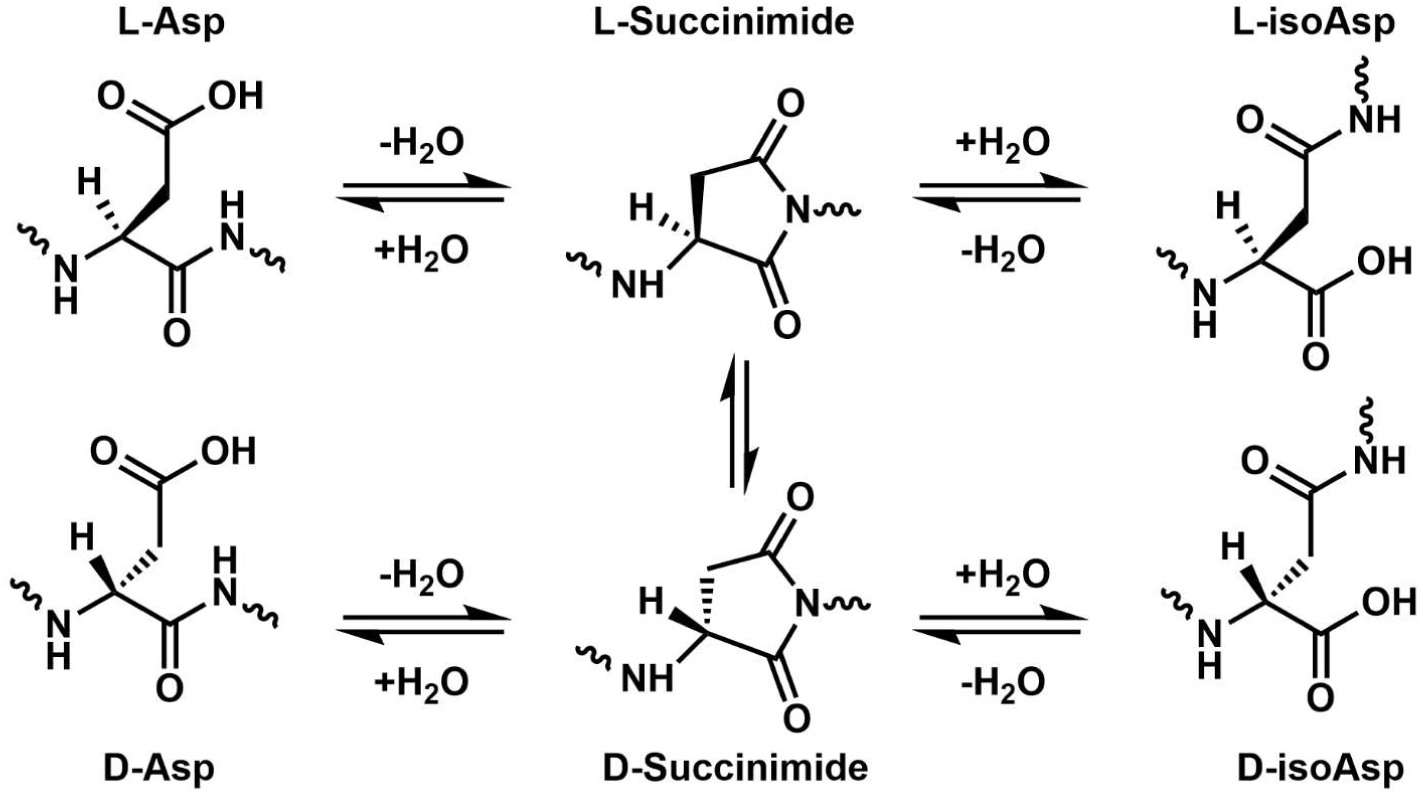
Mechanism of Aspartic Acid Isomerization

Deamidation of asparagine (Asn) also forms the identical succinimide ring intermediate in Scheme 1 and leads to conversion of Asn into all four isomeric forms of Asp.^20^ The addition of methylene into the backbone chains of L-isoAsp and D-isoAsp introduces a “kink” into the peptide backbone, and the sidechain of D-Asp is oriented improperly relative to L-Asp. These changes in 3D conformation have been shown to impact protein function and turnover.^21^ In particular, isomerization impedes the activity of lysosomal cathepsins responsible for maintaining autophagic flux, potentially leading to lysosomal storage.^18,21,22^ Autophagic flux dysfunction and lysosomal failure are phenotypes increasingly associated with neurodegenerative disease.^23–25^ Recently, we have shown that formation of Asp isomers in tau is effectively diagnostic of Alzheimer’s disease, likely due to correlation with a loss of autophagic flux.^26^

The study of isomerization by mass spectrometry (MS) presents a unique challenge because isomers have the same atomic composition and mass. In addition, unique fragment ions are not possible for L- vs D-Asp isomers, and while unique fragments can be generated to distinguish L/D from L/D-isoAsp using electron-based dissociation, the intensity of such fragment ions is often very low or nonexistent.^27–29^ The different 3D structures of the four isomeric forms of Asp allow for identification via differences in fragment ion abundance in tandem-MS,^30–32^ as well as separation by ion mobility^33–35^ and, most commonly, reversed-phase liquid chromatography.^16,18,20,36–38^ However, the dynamic exclusion used in typical data-dependent acquisition data prevents the same mass from being re-analyzed for a chosen period. Given that Asp isomers have the same exact mass, isomer peaks eluting within the dynamic exclusion window will not be examined. In complex biological samples where it is not uncommon for multiple peptides to have similar retention times, a lack of MS^2^ spectra due to dynamic exclusion makes it very difficult to confirm that a given peak corresponds to an isomer. The use of targeted LC-MS approaches such as parallel reaction monitoring (PRM) avoids issues with dynamic exclusion but greatly limits the number of analyzable peptides.^38^ One approach that has shown promise in isomer identification is data-independent acquisition (DIA). By continuously monitoring both precursor and fragment ion signals, DIA affords straightforward confirmation that multiple observed peaks correspond to aspartic acid isomers. The ability to do an untargeted search also allows thousands of potential isomers to be profiled in a single experiment.^26^

While the difficulties in studying isomerization systematically for large numbers of peptides or proteins have limited analysis of isomerization rates, the ∼1 Da mass shift associated with Asn deamidation has facilitated more detailed profiling.^39,40^ Work from the Robinson lab on synthetic peptides suggests that the amino acid C-terminal to Asn strongly influences deamidation,^41,42^ while experiments with proteins have focused on the potential importance of secondary structure.^43–45^ Although deamidation and isomerization lead to the same succinimide ring intermediate and are often assumed to be highly similar processes, potentially important chemical differences between the residues do exist. A few isomerization rates have been experimentally determined for Asp in peptides^19,46^ and in the specific context of antibodies,^47–52^ but overall, the rate of Asp isomerization in intact proteins has received very little attention.

Experiments studying the proteins of the eye lens allow for detection and quantification of isomers across the human lifetime,^8,9,16,37,38,53–57^ but determination of isomerization rates is not possible in such experiments because only a single timepoint can be collected for each individual. Additionally, isomerization rates have not been examined in intact protein complexes or aggregates, both of which have relevance in the context of neurodegenerative disease.

In the present work, we set out to greatly expand the number of isomerization rates determined within intact proteins and protein complexes. Using *in vitro* aging of cell lysates to create isomers in a complex environment, we utilized DIA LC-MS to identify and quantify isomerization for a diverse group of proteins. The potential impacts of primary sequence, secondary structure, protein complex formation, and base catalysis were examined. Our results reveal that multiple levels of protein structure influence Asp isomerization. In particular, steric hindrance by nearby side chains and backbone flexibility impact isomerization. Isomerization is tempered within protein complexes but is not sensitive to small increases in pH. *In vitro* aging of tau revealed faster-than-expected isomerization, which was slowed but not stopped by heparin-induced aggregation.

## Experimental

### Materials

Dulbecco’s phosphate-buffered saline (DPBS) and 10 cm plates were obtained from VWR. Cell detachment solution (0.25% trypsin, 0.1% EDTA, w/v), L-glutamine, penicillin, and streptomycin were obtained from Corning. Fetal bovine serum (FBS) was from Seradigm. Dulbecco’s modified eagle’s medium/Nutrient mixture F-12 Ham (DMEM-F12), Roche protease inhibitor cocktail (PIC) with EDTA, iodoacetamide (IAA), 1,4-dithiothreitol (DTT), tris(2- carboxyethyl)phosphine hydrochloride (TCEP), and 3KDa molecular weight cutoff filters were purchased from Millipore Sigma. Optima-grade water, optima-grade formic acid, urea, sodium chloride (NaCl), sequencing grade trypsin, Tris base, Triton X-100, sodium deoxycholate, magnesium chloride (MgCl_2_), sodium dodecyl sulfate (SDS), and Pierce Peptide Retention Time Calibration Mixture (PRTC) were obtained from Thermo Fisher Scientific. Bradford reagent was purchased from Bio-Rad. 100 µm diameter fused silica was obtained from Polymicro. Reprosil-Pur 120 C18-AQ 3 µm and 1.9 µm were purchased from Dr. Maisch GmbH. Full-length recombinant 2N4R tau protein was purchased from R & D Biosystems.

### Cell Culture

SH-SY5Y cells were obtained from American Type Culture Collection (ATCC) and used within 20 passages. Cells were maintained in DMEM-F12 media supplemented with 1% 2mM glutamine, 1% penicillin, 1% streptomycin, and 10% fetal bovine serum. Cell culture media and cell detachment solution were warmed to 37°C in an aluminum bead bath prior to use. Biological triplicate samples were generated and used in the following experiments.

### Cell lysis

Radio-immunoprecipitation assay (RIPA) buffer was made using 2.5 mL of 1 M Tris pH 7.5, 7.5 mL of 1 M NaCl, 5 mL of 10% w/v Triton X-100, 2.5 mL 10% w/v sodium deoxycholate, 250 µL of 20% SDS, and 32.25 mL of Optima water (final concentrations: 50 mM Tris pH 7.5, 150 mM NaCl, 1% w/v Triton X-100, 0.5% w/v sodium deoxycholate, and 0.1% SDS). Native lysis buffer was made using 2.5 mL 1 M Tris pH 7.5, 2.5 mL 1 M NaCl, 5 mL 100 mM MgCl_2_, 1.5 mL 10% w/v Triton X-100, 500 µL of 100 mM DTT, and 33 mL Optima water (final concentrations: 50 mM Tris pH 7.5, 150 mM NaCl, 10 mM MgCl_2_, 0.3% w/v Triton X-100, and 1 mM DTT). Both buffers were prepared alongside variants where Tris pH 7.5 was adjusted to pH 8.8 with ammonium hydroxide. Cells were grown to confluence, washed with DPBS, harvested in DPBS by scraping, and lysed on ice for 30 minutes with lysis buffer containing protease inhibitor cocktail. Lysates were clarified by centrifugation at 21,100 g for 15 minutes at 4°C. Soluble protein concentration was determined by colorimetric assay (Bio-Rad) using Agilent Cary 60 UV-Vis spectrophotometer, and lysate concentration was normalized to 1 µg/µL with 50 mM Tris at appropriate pH.

### *In vitro* aging and sample cleanup

Lysates were split into 5 low-binding microcentrifuge tubes with 100 µL of lysate each, one for each time point, tightly covered with Parafilm, and placed into a 37°C sand bath. At each chosen time point, (Days 0, 5, 10, 25, and 50), samples were removed from the sand bath. Only MS-grade organic solvents were used during sample cleanup and preparation, excluding chloroform (CHCl_3_). Sequentially, 400 µL MeOH, 100 µL CHCl_3_, and 300 µL H_2_O were added and gently vortexed. After centrifugation twice at 21,100 G for 5 minutes, protein pellets formed between the interface, and the majority of the top layer was removed by aspiration. The remaining solution was cleaned by addition of 400 µL MeOH, vortexing, centrifugation at 21,100 G, and supernatant aspiration, repeated ≥5 times. Pellets were air-dried gently overnight.

### Sample digestion and preparation

Dried protein pellets were resuspended in 100 µL of 8M urea and stored at 4°C for 24 hours to fully dissolve the pellet. 10 µg of protein (10 µL of solution) was transferred to a low-binding microcentrifuge tube, and 200 ng of yeast enolase (1 µL of solution) was added as a process control. Samples were then reduced by addition of 2 µL of 25 mM TCEP for 20 minutes at ambient temperature, alkylated by addition of 2 µL of 50 mM IAA for 60 minutes in the dark at ambient temperature, diluted with 25 µL of Tris buffer to 2 M urea, and digested with 0.2 µg trypsin (1:50) overnight at 37°C for 16-20 hours. Tryptic digestion was quenched by adding 2 µL of formic acid to pH 2.0 (verified by pH paper). Samples were filtered using 0.2 µm filters to remove any particulates and then stored in the freezer at ≤ -60°C until analysis.

### Column preparation

Monophasic C18 analytical columns were prepared by pulling 100 µm diameter fused silica columns with a P-2000 laser tip puller (Sutter Instrument Co., Novato, CA) and packing to a length of 30 cm with Reprosil-Pur 120 C18-AQ 3 µm resin.

### Data-independent acquisition mass spectrometry

200 ng of each sample with 250 femtomole of PRTC was analyzed using a Thermo UltiMate 3000 RSLC interfaced to a Thermo Fisher Scientific Orbitrap Fusion Lumos Tribrid Mass Spectrometer. PRTC was added to act as a retention time calibrant and assess the performance of the instrument during acquisition.

Samples were eluted using 0.1% formic acid in water as mobile phase A and 0.1% formic acid in 80% acetonitrile as mobile phase B. Samples were desalted by loading onto an Acclaim PepMap 100 C18 3 µm 75 µm x 2 cm trap cartridge at 1% B for 1.8 minutes. Peptides were then separated with a 90-minute LC gradient consisting of 1% B for 3 min, 1-7% B in 7 min, 7- 15% B in 20 min, 15-50% B in 40 min, 50-98% B in 5 min, 98% B for 10 min, and a wash of 1% B for 5 min. The analytical column was then equilibrated at 1% B for 100 minutes. For all experiments, the mass spectrometer was operated in data-independent acquisition mode. Each run consisted of a cycle of one 60,000 resolution SIM scan mass spectrum with a mass range of 330-1500 m/z and an automatically determined AGC target followed by data-independent MS^2^ spectra on the loop count of 71 at 15,000 resolution, an automatically determined max injection time of 22 ms, a normalized AGC target of 1000%, and 33% HCD collision energy with an overlapping 8 m/z isolation window.^58–60^ A total of 60 LC experiments were conducted (5 time points with 4 lysis buffer conditions and biological triplicates).

### Data Analysis

Thermo Raw files were converted into the mzML format using Proteowizard MSConvert (version 3.0.22288) using vendor peak picking and demultiplexing settings.^61,62^ A quantitative spectral library was generated using default settings in EncyclopeDIA (version 2.12.30)^63–65^ by using a Prosit predicted spectral library^66^ and all 60 LC runs, requiring a minimum of three quantitative ions and filtering peptides at 1% FDR using Percolator (version 3-01). ^67,68^ The quantitative spectral library was then imported into Skyline (daily version 23.1.1.268) with the human Uniprot FASTA with yeast enolase added as the background proteome to map peptides to proteins.^69,70^ Transition retention time filtering settings were set to “use only scans within 5 min of MS/MS IDs” for all runs.

### Isomer identification and quantification

Isomers were manually identified in the dataset by the presence of two or more chromatographic peaks with the same fragments observed at different retention times. Since asparagine deamidation occurs on a relevant timescale to these experiments and may generate multiple chromatographic peaks, asparagine-containing peptides were removed from the search by filtering with Skyline’s document grid. Isomers containing multiple, non-sequential aspartic acid residues were also removed since it would be unclear which Asp contributed to which chromatographic peaks. Sequence isomers, which also generate multiple chromatographic peaks due to different amino acid order, were identified based on the presence of multiple peaks at Day 0 and were removed from downstream analysis (see Figure S1 for an example). Isomer quantification was performed as described previously^26^. Briefly, peak values were extracted from Skyline by manually adjusting the boundaries for each individual isomer peak. Only fragment ion areas were used for quantification to limit the impact of precursor interference. The total areas of the fragment peaks corresponding to the peptide of interest were summed to produce a “total isomer area”. Although fragmentation yield has been shown to vary between isomers, the overall fragmentation efficiency does not change regardless of changes to individual fragments.^31^ To calculate percent isomerization values, the “total isomer area” was divided by the total peak area including the non-isomer peak. Means and standard deviation values were calculated in Microsoft Excel using the three biological triplicate values for each peptide’s percent isomerization at each time point. Rate constants for isomerization were calculated assuming that isomerization is a first-order reaction up through Day 50. When comparing between two buffer conditions, peptides were required to show some degree of isomerization in both conditions and not originate from proteins that were expected to be differentially extracted, such as membrane proteins being excluded since they would not be well extracted by Native lysis buffers. All figures were generated using OriginPro 2024 (10.1.0.178). Scheme 2 was generated using BioRender.

**Scheme 2.**
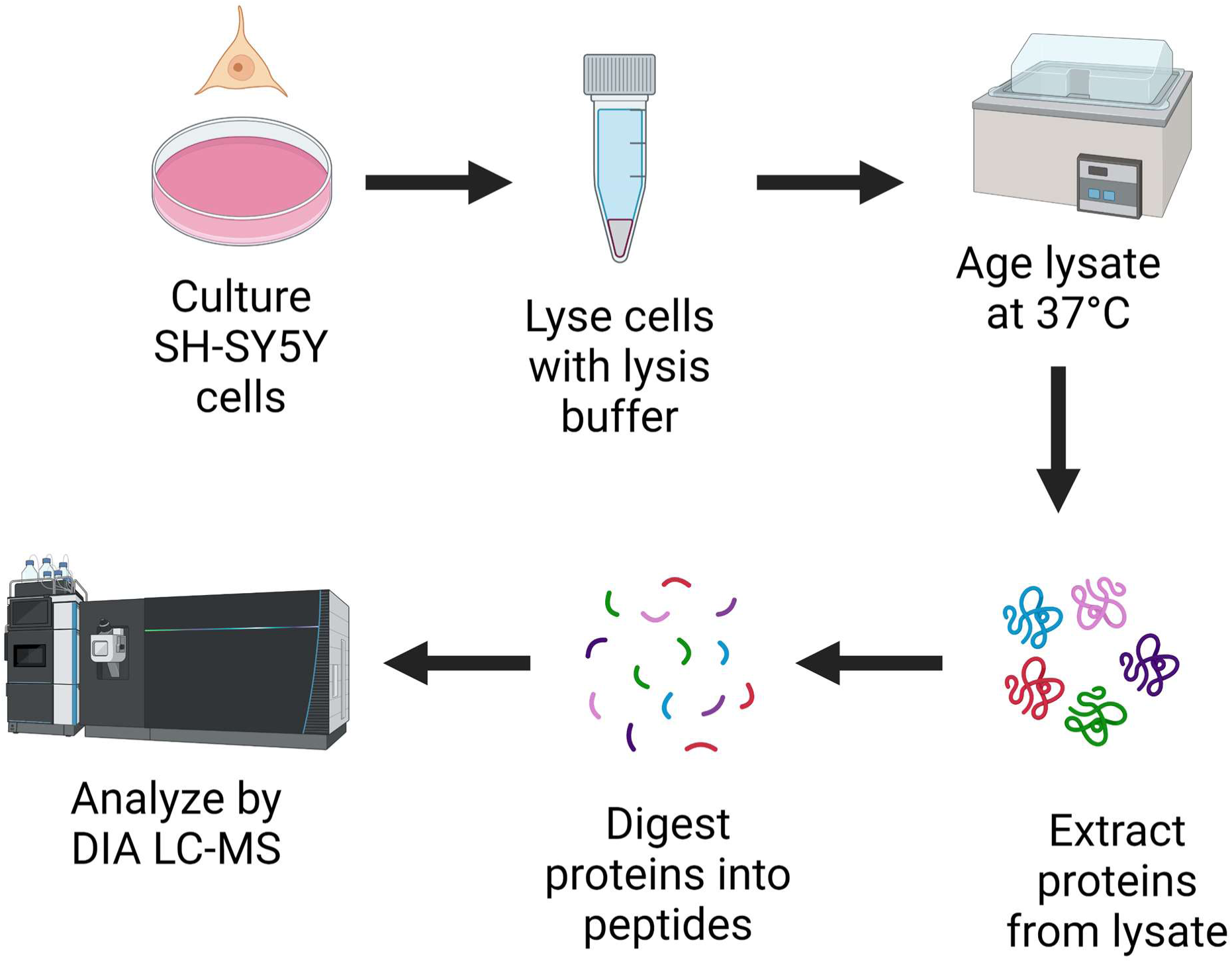
Experimental procedure for *in vitro* aging of cell lysates and preparation for LC-MS

### Analysis of protein structure

PDB files for each protein (see Table S1) containing a quantifiable isomer were identified in Uniprot and imported into Maestro (version 13.4). When crystal structures, EM structures, or NMR structures containing the aspartic acid of interest were unavailable, AlphaFold structures were used. To quantify the amount of crowding around the C-terminal amide that forms the succinimide ring intermediate, the number of atoms, excluding hydrogens, within 4 Å of the nitrogen atom were counted using the “Display all distances” tool.

Hydrogen bonding interactions were also recorded as present or absent at the hydrogen atom in the C-terminal amide. The secondary structure of the aspartic acid residue of interest was classified as being located on an alpha helix, a beta sheet, or a non-regular region, such as turns, where the structures have nonrepeating backbone torsion angles.^71^

### Identification and quantification of Asn deamidation

Deamidation sites often generate several deamidated peaks due to the formation of a succinimide ring intermediate and subsequent Asp isomers, so deamidation sites were identified similarly to Asp isomerization by the presence of multiple chromatographic peaks. Due to the ∼1 Da mass shift associated with deamidation, loss of fragment ion intensity for fragments containing the Asn residue can occur depending on the fragments generated. This aids in confirmation that a peptide is deamidated and helps localize the deamidation site. For quantification, only fragment ions found in both non-deamidated and deamidated peaks were used, with a minimum of 3 fragment ions being required for a peptide to be considered quantifiable. %Deamidation values for Asn sites were calculated by the same method as percent isomerization for Asp isomers. All Asp-containing peptides were excluded when searching for deamidation to remove interference.

### *in vitro* aging of recombinant tau

Non-aggregated tau samples were aliquoted in 1X PBS buffer, pH 7.4. Each aliquot was 50 µL with a concentration of 4.35 µM. Aggregated samples were prepared using heparin sodium salt. Aliquots of 35 uL of 10 µM protein in 1X PBS, pH 7.4 with 1 mM DTT, 2 mM MgCl_2_, and approximately 10 µM heparin sodium salt were shaken for 48 hours at 37°C to induce aggregation.^72^

Samples were placed in 500 µL Eppendorf tubes and placed inside glass vials with 1 mL of water to prevent solvent evaporation. Samples were incubated in a sand bath at 37°C. Timepoints were taken at 0, 3, 6, 9, 12, and 16 weeks. After collection of the timepoints, they were treated with 50 µL of 70% formic acid at room temperature for two hours in order to monomerize any aggregated protein. All solvent was then removed by lyophilization, and samples were resuspended in 10 µL of 50 mM tris-HCl pH 8.0 with 6 M urea.

### Tau digestion

Cysteines were reduced and alkylated by adding 3 µL of 10 mM dithiothreitol and incubated 20 minutes at room temperature followed by adding 15 µL of 20 mM iodoacetamide and incubated at room temperature in the dark for 1 hour. Following cysteine alkylation, 100 µL of 50 mM Tris, pH 7.6 was added (urea concentration ≤0.6 M). The pH was checked, and if necessary adjusted to pH 8-9 using 1 M Tris-HCl, pH 9. Trypsin was added in a 1:20 enzyme:protein ratio and incubated for 18 hours at 37°C. Samples were then filtered and 100 µL of optima water with 0.1% formic acid was added. Aggregated samples were also run through a 3KDa molecular weight cutoff filter and the flow through was retained to remove any large irreversible aggregates.

### LC-MS/MS Analysis

Samples were analyzed using the same instrument as above. LC separation was performed using a 25 cm 100 µm column packed in house with 1.9 µm C18-AQ resin. Mobile phase A was Optima water with 0.1% formic acid, and mobile phase B was 80% acetonitrile, 20% Optima water, and 0.1% formic acid. The gradient was 1-7% B in 7 minutes, 7- 15% B in 20 minutes, 15-50% B in 40 minutes, and 50-98% B in 5 minutes. A parallel reaction monitoring (PRM) method was used with 60 targets. The MS1 resolution was 120,000 with a scan range of 350-1500 m/z. MS2 used HCD with NCE 33, an isolation window of 1.6, resolution of 15,000, and scan range of 350-2000 m/z.

## Results and Discussion

### Detection of isomerization in cell lysates by DIA

To simultaneously monitor the isomerization of many intact proteins, whole cell lysates from SH-SY5Y cells were utilized. For one set of experiments, RIPA buffer was selected for cell lysis to efficiently solubilize monomeric proteins while disrupting protein-protein interactions.^73^ Due to the high protein turnover rate in replicating cells, freshly prepared lysates from SH-SY5Ys were predicted to contain few Asp isomers. To induce isomerization, cell lysates were incubated for varying lengths of time at 37°C prior to protein extraction, digestion, and analysis by DIA-LC-MS as summarized in Scheme 2.

A representative example of results illustrating the progression of isomerization over time is shown in Figure 1. More specifically, the intensities of fragment ions used to identify the Asp-containing peptide ALVDGPCTQVR are plotted as a function of elution time for 5 incubation periods. On Day 0, a single chromatographic peak cluster is present, and importantly, all the colored lines representing different fragment ions are well aligned. After 5 days of aging, a second peak cluster composed of the same fragment ions appears at a earlier retention time. By Day 50, four chromatographic peak clusters are observed for the peptide, each representing a different isomer of Asp. Based on their relative retention times, isomer peaks are clearly differentiable from the L-Asp peak. Additionally, close examination of the L-Asp and Isomer 2 peaks reveals that the relative abundance of the fragment ions y_7_ and y_8_ invert, as well as y_6_ and y_9_. Drawing on previous work, observing changes in the relative abundance of fragment ions in tandem MS is a frequently observed phenomenon for peptide isomers.^30,31^ Furthermore, increasing peak intensity over time and the presence of all fragment ions in each peak rules out other types of isomers that may separate by liquid chromatography, such as sequence isomers (see Figure S1 for an example sequence isomer). These chromatograms clearly show that while Asp isomers are not present in SH-SY5Y cell lysates originally, the passage of time allows all four isomeric forms of Asp to be generated and observed.

**Figure 1.**
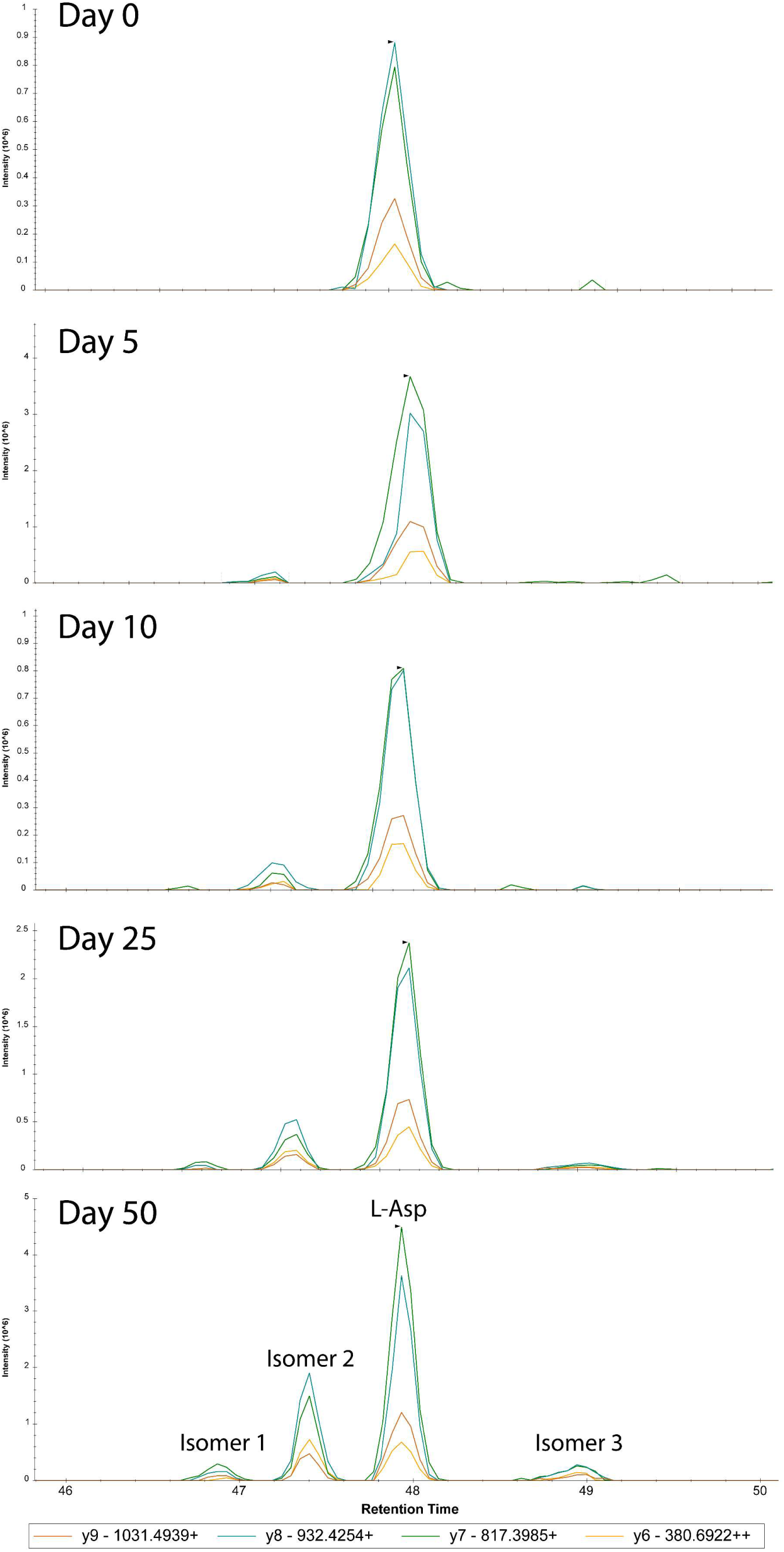
Skyline chromatograms for the peptide ALVDGPCTQVR from the protein RL14 across 5 *in-vitro* aging time points. Each colored line corresponds a fragment ion associated with the precursor in the spectral library. The first isomer peak emerges at a different retention time on day 5, with all four peaks being clearly visible by Day 50.

### Influence of primary sequence

Prior work by the Robinson group has shown that deamidation of Asn within a peptide is most strongly influenced by the residue C-terminal to it.^41,42^ Bulkier amino acids, such as Trp or Tyr, slow deamidation while C-terminal Gly yielded the highest rates. These results were rationalized in terms of steric hindrance and crowding of the reaction site, as illustrated in Scheme 3 for Asn-Gly and Asn-Phe with the reaction site highlighted in yellow. For Asn-Gly, the reaction site is open regardless of the conformation adopted by the peptide, while for Asn-Phe, some conformations will lead to crowding of the reaction site by the Phe sidechain. For small, presumably disordered, peptides such as those used by the Robinson group, many conformations will be sampled over time. Thus, the differences in deamidation rate observed by the Robinson group likely reflect an average steric hindrance, with bulkier side chains being more likely to hinder the reaction site.

**Scheme 3.**
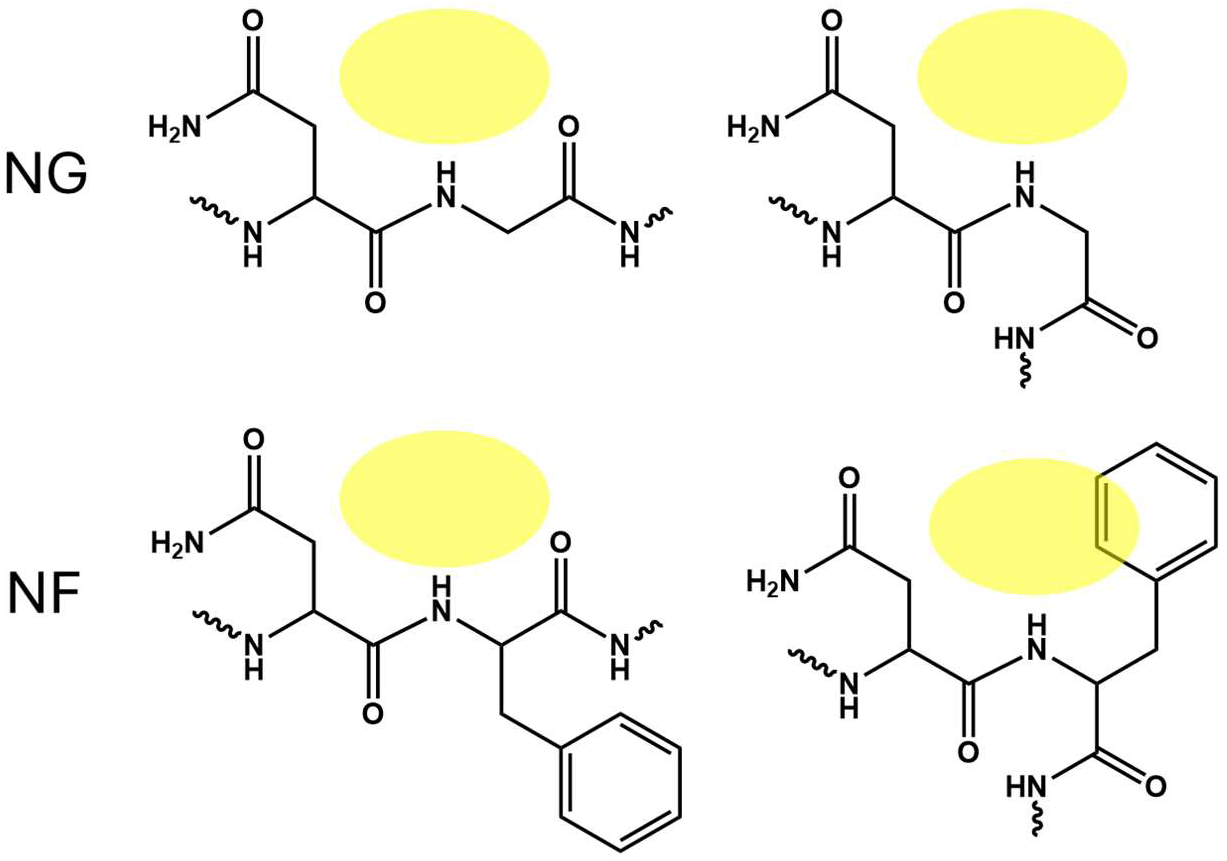
Examples of potential conformations for deamidation site residue pairs, with the reaction site allowing succinimide ring formation highlighted in yellow.

Given that Asn deamidation leads to the same succinimide ring intermediate as Asp isomerization, we investigated the relationship between primary sequence and Asp isomerization rate in our data. Figure 2a shows the distribution of Asp isomers binned by the C-terminal amino acid residue. Just over 50% of all observed isomers occur at DG sites, with DS and DA sites yielding the next highest frequencies. These observations are consistent with steric hindrance being a primary influence of Asp isomerization. To confirm that our observations are not overly influenced by the inherent distribution of each residue pair in the proteome, we determined the number of peptides present for each DX site. Figure S2 shows that DG sites are not heavily enriched, and more common sites such as DL do not yield large numbers of isomers.

**Figure 2.**
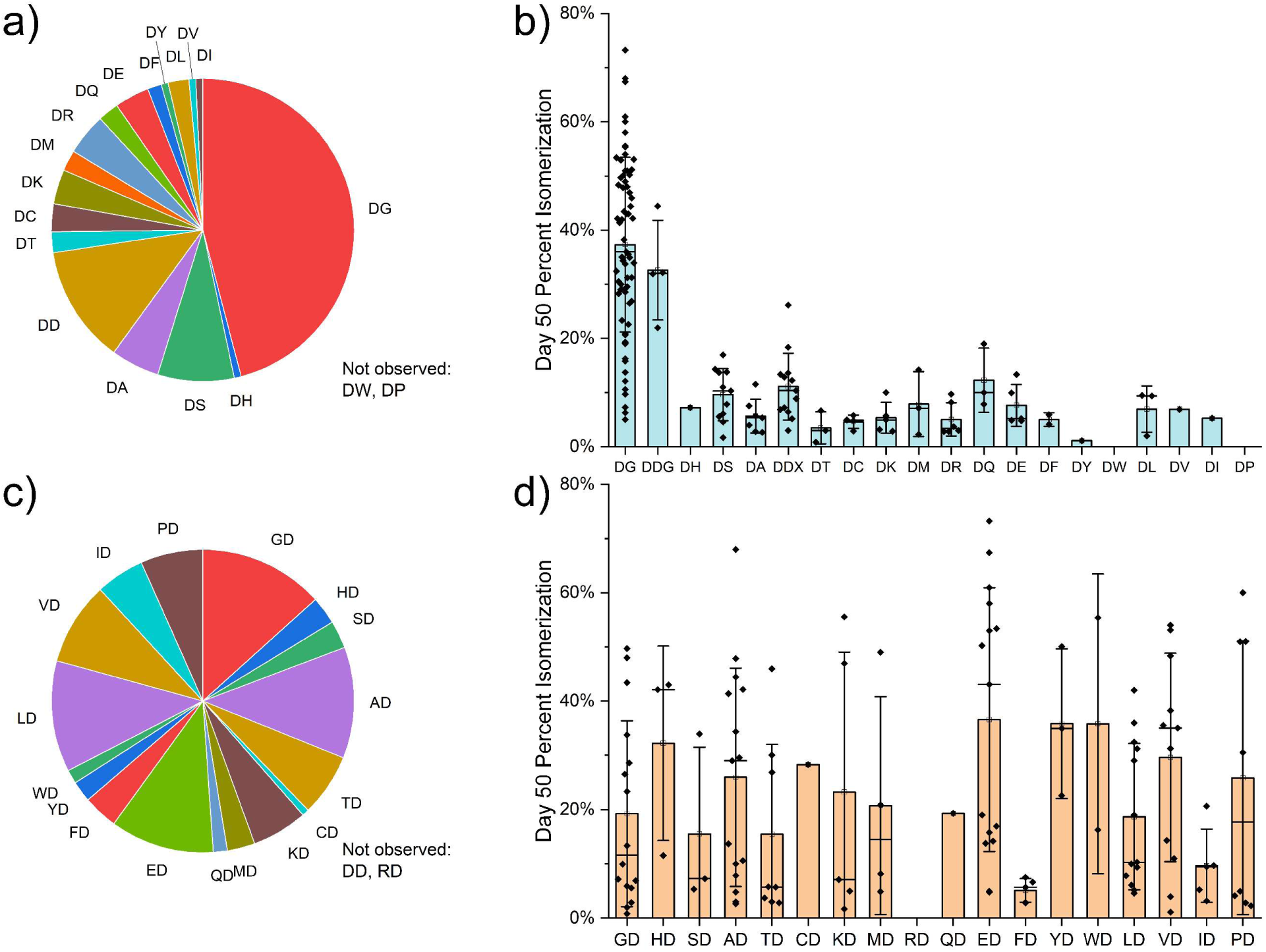
Impact of primary sequence on isomerization. a) Number of detected isomers and b) percent isomerization binned by the residue C-terminal to Asp. c) Number of detected isomers and d) percent isomerization binned by the residue N-terminal to Asp. Error bars for b) and d) represent one standard deviation.

The amount of isomerization after 50 days of aging is shown in Figure 2b, again sorted by C-terminal residue. In addition to occurring with greater frequency, DG isomers often undergo greater isomerization than non-DG sites. The range of DG isomerization is significantly larger than that observed for non-DG sites, which tend to cluster around 10%. For DD isomers, it is unclear which Asp residue isomerizes, so these peptides are grouped as DDG isomers, which isomerize quickly, and DDX isomers, which reach similar isomerization levels to all other motifs. Additional clarity for the importance of the C-terminal residue can be obtained by performing the same analysis for the N-terminal residue, as shown in Fig. 2c and 2d. There are no clear trends in the frequency or extent of isomerization based on the N-terminal amino acid identity, consistent with observations for deamidation.^41^ Taken as a whole, the trends observed in isomerization rate match well with prior work on deamidation, and it is clear that even in the highly complex environment of intact proteins, steric hindrance by the C-terminal amino acid side chain is a major determinant of isomerization rate.

### Crowding of the C-terminal amide

Although Figure 2 clearly shows that the identity of the C-terminal amino acid plays a major role in determining the isomerization rate for a given Asp residue, the wide range of values for DG isomers suggests that other factors also play a role. In contrast to short, unstructured peptides, the orientations of the side chains of amino acids C-terminal to Asp are determined by secondary and tertiary structure in intact proteins. Alpha helical and beta sheet structures have consistent orientations of the constituent side chains, while non-regular structures, such as turns, yield inconsistent peptide backbone torsion angles and sidechain orientations.^74^ Additionally, while steric hindrance in proteins may be due to the side chain of the C-terminal amino acid, crowding could also be caused by sequence-remote interactions. To quantify the degree of crowding caused by any interacting component in the three-dimensional structure for each protein, all atoms within 4 Å of the amide nitrogen C-terminal to any isomerized Asp were counted. As shown in Figure 3a, there is a negative correlation between percent isomerization and crowding, implying that as the amide becomes more crowded, steric hindrance increasingly limits formation of the succinimide ring. To separate the impact of crowding by the C-terminal side chain and crowding from sequence-distant interactions, Figure 3b shows the same results sorted by DG and non-DG peptides.

**Figure 3.**
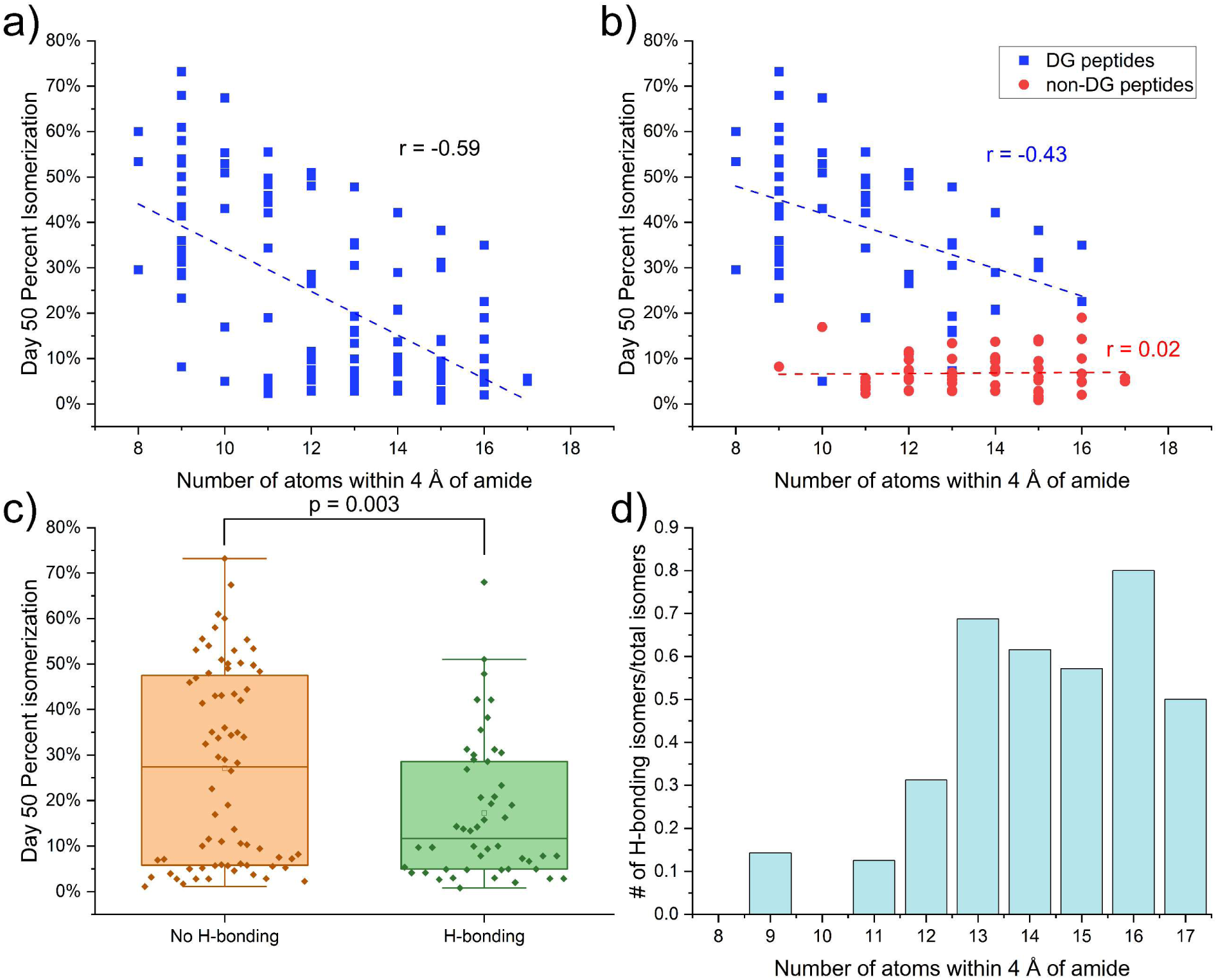
Crowding of the C-terminal amide. (a) percent isomerization compared to the number of atoms within 4 Å of the C-terminal amide involved in succinimide formation (b) percent isomerization compared to the number of atoms within 4 Å of the C-terminal amide involved in succinimide formation, binned by DG or non-DG isomer. (c) Box-and-whisker plot showing percent isomerization binned by presence of absence of hydrogen bonding (d) Ratio of the number of isomers where the hydrogen of the C-terminal amide has hydrogen bonding interactions to the total number of isomers.

Isomers with non-DG sites tend to have both increased amounts of crowding and lower percent isomerization than DG sites, suggesting that the number of atoms within 4 Å of the amide primarily results from the presence of a side chain on the C-terminal residue, and that longer-range interactions are less important. However, for the DG peptides, there is still a moderate, negative correlation between crowding and percent isomerization. The atoms leading to increased crowding in DG sites could originate from either the orientation of amino acids close in sequence or sequence-distant interactions due to protein tertiary structure. As shown in Figure S3a, in one case of high crowding, an N-terminal tyrosine led to significantly increased steric hindrance. However, Figure S3b shows an alternate situation where atoms were localized towards the amide from ∼20 amino acids away in sequence, a long-range interaction. These results demonstrate that in addition to residues nearby to the C-terminal amide, tertiary protein structure can increase crowding and impact isomerization propensity.

Hydrogen bonding interactions represent another type of sequence-distant interaction that is crucial to the structure and function of proteins.^75–78^ Given the strength of hydrogen bonding interactions, we hypothesized that hydrogen bonding to the C-terminal amide could make steric clashes more persistent. To evaluate this possibility, we checked each C-terminal amide for hydrogen bonds. Figure 3c shows percent isomerization binned by the presence or absence of hydrogen bonds. On average, isomerization is lower in the presence of hydrogen bonds. This suggests that hydrogen bonding interactions may lead to increased crowding, so to examine this more closely, Figure 3d shows the ratio between the number of isomers that form hydrogen bonds to the total number of isomers. More crowded isomers tend to also have more hydrogen bonding interactions, which could be due to atoms involved in the hydrogen bond being localized towards the amide. Interestingly, the proportion of hydrogen bonded isomers is relatively stable after 13 nearby atoms. Given that hydrogen bonded isomers have lower isomerization and atoms from hydrogen bonding partners can only contribute so much to crowding, it is likely that H-bonds make less crowded interactions more persistent. This persistence can lower isomerization rates independent of the amount of crowding. Overall, the rate of isomerization in intact proteins is clearly influenced by crowding of the C-terminal amide, both by side chain atoms and sequence-distant residues, as well as the persistence of interactions due to hydrogen bonding.

### Impact of secondary structure

In addition to the steric environment of the C-terminal amide, previous work suggests that backbone flexibility may also influence succinimide ring formation. Kosky et. al. showed that deamidation in peptides is greatly limited by increasing helicity, suggesting that alpha helical structure restricts the flexibility required to form the succinimide ring intermediate.^79^ Work from the Robinson lab similarly found that deamidation is slowed by alpha helical regions within intact proteins.^43^ To determine whether backbone rigidity influences Asp isomerization, isomer sites were classified based on their secondary structure as alpha helical, beta sheet, or non-regular. Figure 4a shows that most isomers are located in non-regular regions, with a smaller subset found in alpha helices. Notably, very few isomers are located in beta sheets. These results may imply that while both disfavor isomerization, alpha helices are more easily distorted to accommodate succinimide ring formation compared with beta sheets, which is consistent with the respective inherent stabilities.^80^ To examine whether the extent of isomerization also depends on secondary structure, percent isomerization values were plotted for isomers binned by structure and C-terminal residue, as shown in Figure 4b. For non-DG isomers, the extent of isomerization is similar regardless of the underlying secondary structure. However, isomerization of DG isomers is highest in non-regular regions, notably less for alpha helices, and very low for beta sheets (albeit for a single data point). Figure 4c illustrates the relationship between crowding and secondary structure. Isomers located in alpha helical sections are often crowded compared to isomers in non-regular regions, suggesting that formation of alpha helices orients side chains close to the C-terminal amide. However, all observed beta sheet isomers are not located in crowded regions, suggesting that backbone rigidity or higher stability must prevent isomerization in the absence of crowding. Taken as a whole, the frequency and extent of isomerization is often decreased in highly structured regions, including both beta sheets and alpha helices, and implicates both crowding and backbone rigidity as factors that influence isomerization.

**Figure 4.**
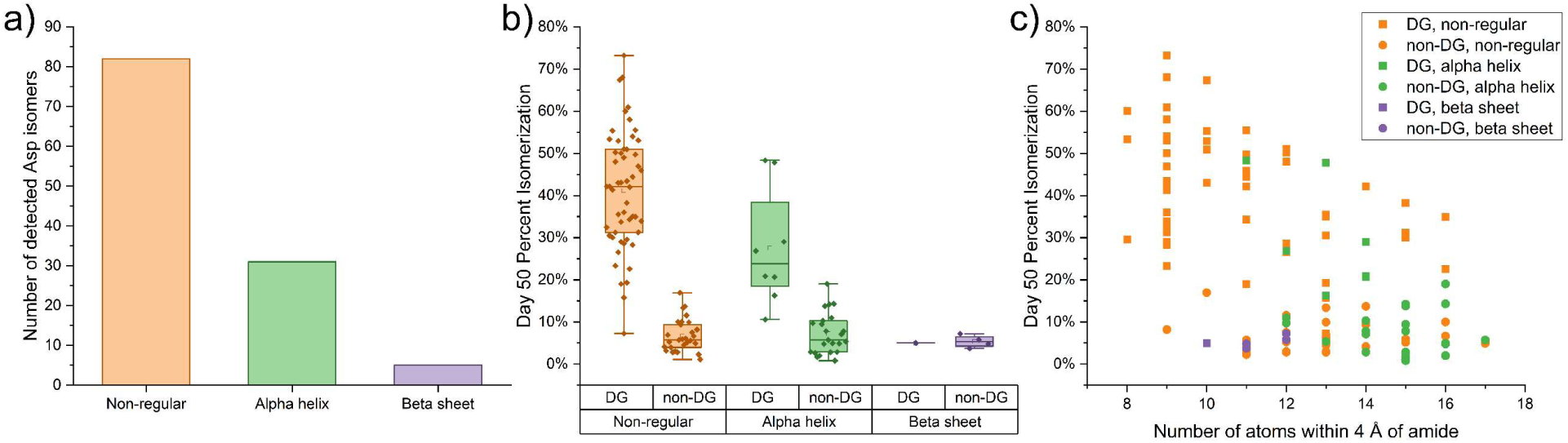
Impact of secondary structure on isomerization. (a) Number of detected Asp isomers sorted by secondary structure element (b) percent isomerization binned by both primary sequence and secondary structure element (c) percent isomerization compared to the number of atoms within 4 Å of the C-terminal amide involved in succinimide formation, binned by secondary structure element.

### Protein-protein interactions

The formation of complexes is a widespread phenomenon that often alters protein folding dynamics and is necessary for function.^81,82^ To investigate the influence of protein-protein interactions on isomerization, we prepared and aged cell lysates using only Triton as a surfactant (termed Native buffer), which was found by Mei et. al. to preserve protein complexes during cell lysis.^83^ Figure 5a shows the progression of isomerization over time for RIPA and Native lysis buffers for the peptide ATAVVDGAFK from PRDX2.

**Figure 5.**
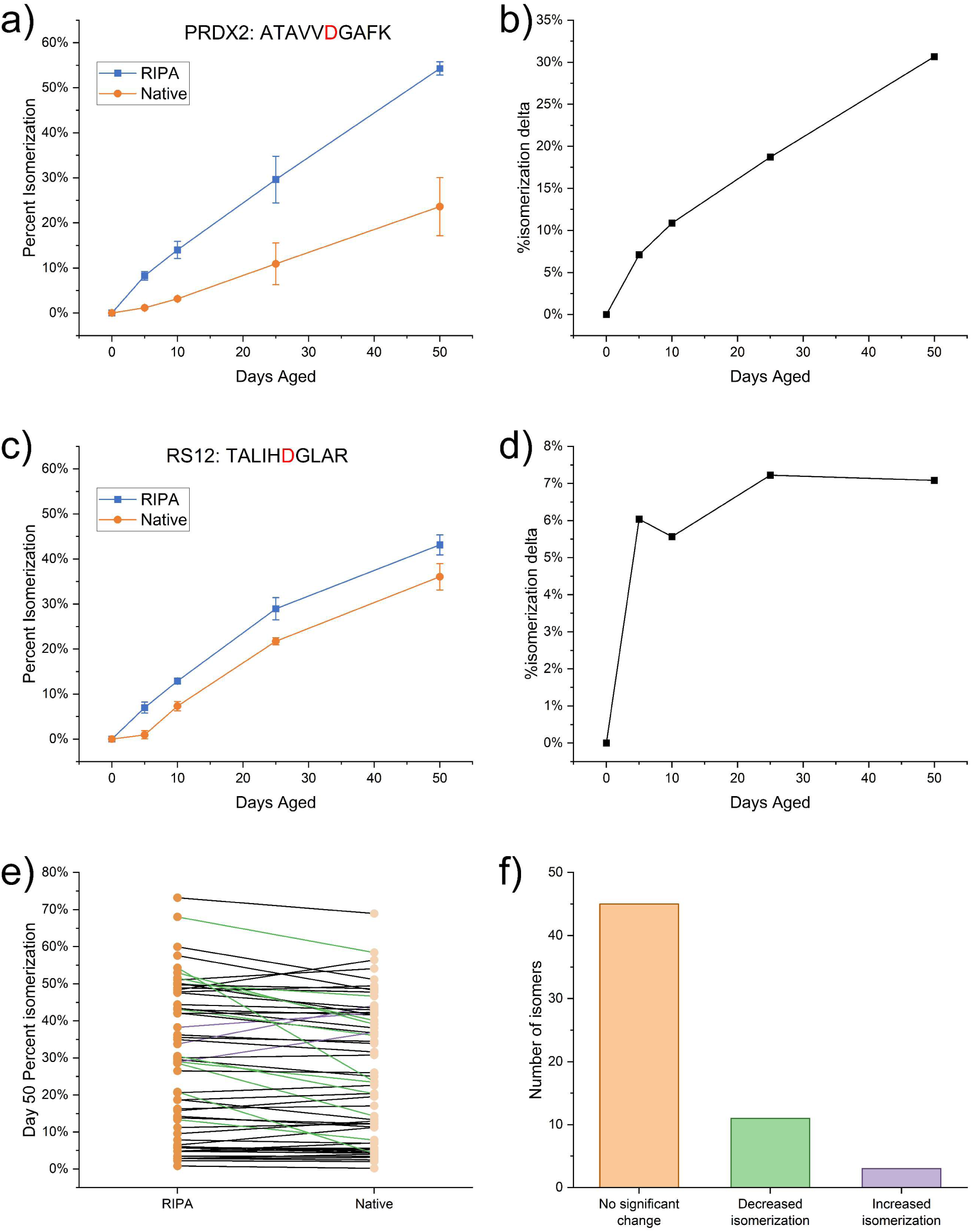
Impact of Native lysis buffer on isomerization. (a) Isomerization at each timepoint in RIPA and Native buffers for the peptide ATAVVDGAFK from PRDX2, error bars correspond to biological replicates (n=3). (b) Difference in percent isomerization (delta) across timepoints for ATAVVDGAFK. (c) Isomerization at each timepoint in RIPA and Native buffers for TALIHDGLAR from RS12, error bars correspond to biological replicates (n=3). (d) Difference in percent isomerization (delta) across timepoints for TALIHDGLAR (e) Before-and-after plot comparing percent isomerization at Day 50 in RIPA and Native lysis buffers for isomerized peptides. Green and purple lines signify decreases and increases, respectively, in percent isomerization greater than two standard deviations of the biological triplicate error. (f) Number of isomers with significant changes to percent isomerization value when comparing RIPA and Native lysis buffers.

Isomerization in Native lysis buffer is clearly reduced throughout the entirety of aging, suggesting that in the intact protein complex, the rate of isomerization is slowed. To quantify the impact on isomerization, Figure 5b shows the difference between the percent isomerization in Native and RIPA lysis buffers for each time point. The difference increases across all timepoints, suggesting that protein-protein interactions are preserved and slow isomerization throughout the entire aging period. PRDX2 forms a homo-decamer ring,^84^ and these results suggest that this complex is highly stable under aging stress.

Another possible outcome is illustrated in Fig. 5c and 5d for the peptide TALIHDGLAR from the protein RS12. While isomerization in Native and RIPA buffers differs at each timepoint, the percent isomerization delta primarily emerges between Day 0 and Day 5, with subsequent isomerization occurring at similar rates. This suggests that the protein complex remained intact initially and decreased isomerization, but after the first 5 days, the complex dissociated, and isomerization was no longer slowed. RS12 is a known binding partner in the 80S ribosome complex^85^, but it did not appear to remain bound during our aging period. Collectively, these results suggest that isomerization could be used to measure and track protein complex stability.

Figure 5e shows a comparison between percent isomerization at Day 50 for all isomers in both RIPA and Native lysis buffers. Significant differences in percent isomerization, labeled with red lines, indicate that the difference between conditions was greater than two standard deviations of the biological triplicate results. The results show that while significant differences are observed, altered isomerization in the Native buffer is relatively uncommon. Figure 5f tabulates the number of isomers by difference in isomerization. Most do not change between buffers, with some decreasing in isomerization under Native buffer, and a few increasing.

Inspection reveals that all the peptides that differ in isomerization propensity between buffer conditions are located within proteins that form protein complexes or promiscuously interact with other proteins (see Table S2). Also shown in Table S2 is a comparison of crowding for each protein in the monomeric and protein complex state, for which no differences were observed.

Illustrative examples are provided in Figure S4, showing that isomer sites need not be located near other subunits for isomerization to be hindered. Taken together, these results imply that global changes in rigidity can be induced by the formation of protein complexes and lead to subsequent alterations in isomerization rate and propensity.

### Impact of basicity on isomerization

Previous work from our lab has shown that the addition of ammonia to a peptide during Asn deamidation leads to an increase in the deamidation rate without significantly altering the relative populations of Asp isomers.^86^ Work from Tyler-Cross and Schrirch suggested a mechanism for general base catalysis by ammonia, where the base assists in the removal of the Gly amide proton during formation of the succinimide ring.^87^ However, Oliyai and Borchardt showed that Asp isomerization in short peptides was not accelerated by ammonia.^88^ To evaluate the influence of pH on Asp isomerization in proteins, we prepared lysates using “accelerated” RIPA and Native buffers where ammonium hydroxide was used to shift the pH from 7.8 to 8.8. Figure 6a shows that isomerization did not change for most peptides in either RIPA or Native buffers at higher pH. In Figure 6b, it is clear that for the small number of peptides affected by pH, isomerization was equally likely to be increased or decreased. To confirm that our observations could not be attributed to an unforeseen experimental flaw, we also examined the effect on deamidation, as shown in Figs. 5c and 5d. Deamidation clearly increases with higher pH, as expected. These results show that isomerization of Asp is not sensitive to minor changes in pH, even for intact proteins and protein complexes, and any changes in isomerization are likely owed to pH-dependent structural changes. In addition, it is clear that given their substantial differences in response to pH, Asp isomerization should not be equated with Asn deamidation.

**Figure 6.**
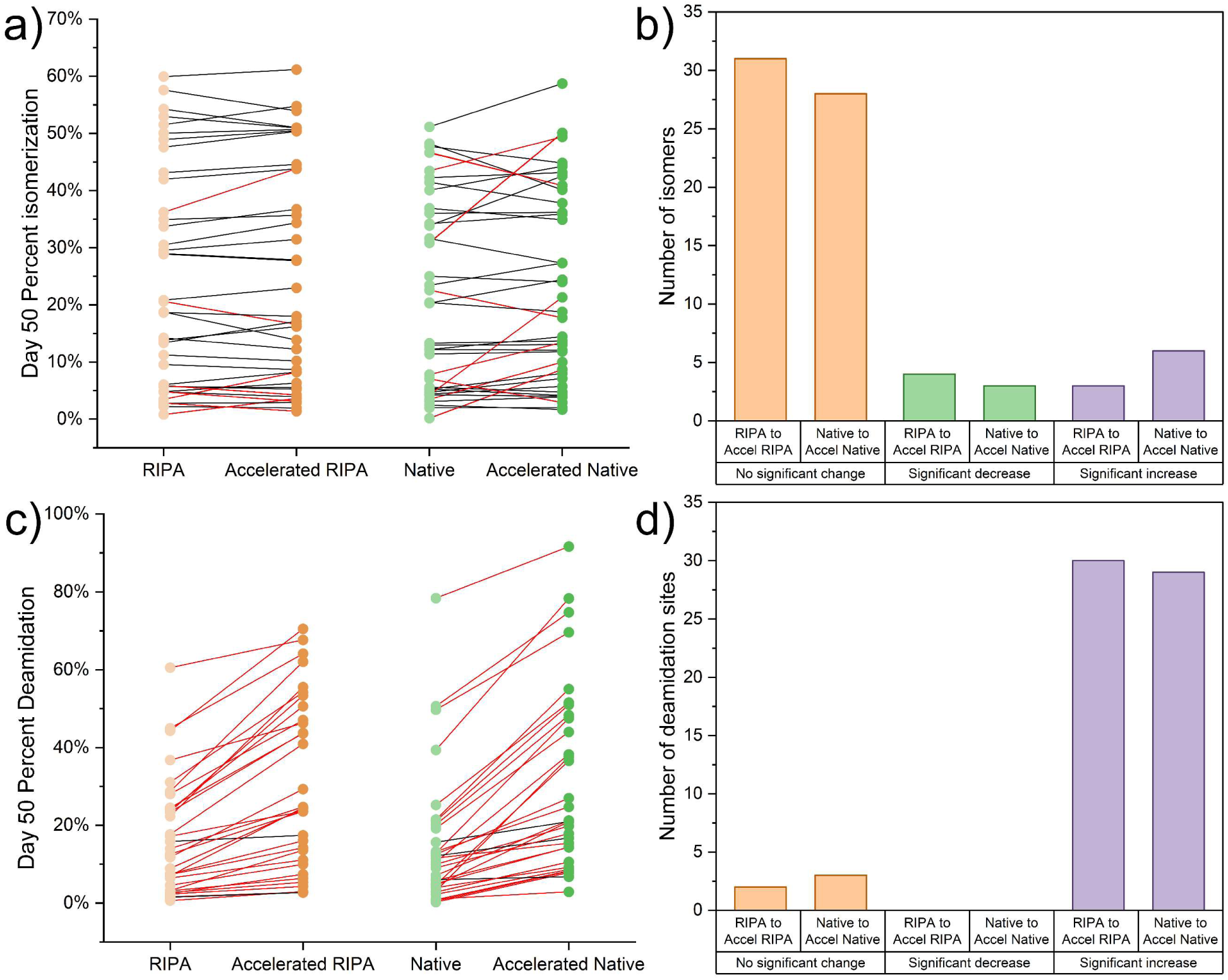
Evaluation of base catalysis and effects of pH on Asp isomerization. (a) Comparisons between “Accelerated” and standard lysis buffers for RIPA (orange) and Native (green) lysis buffers, where red lines correspond to changes in percent isomerization with a difference greater than two standard deviations from triplicate percent isomerization calculations. (b) Bar plot showing whether observed differences led to no significant change (orange), significant decreases (green), or significant increases (purple) for both RIPA and Native lysis buffers. (c-d) Impact of ammonium hydroxide on Asn deamidation at Day 50, shown as in (a) and (b).

### Isomerization of Tau

Having established the primary factors that influence isomerization in proteins, we next sought to use this information to provide greater context for understanding isomerization in aging and neurodegeneration. Tau is strongly associated with Alzheimer’s disease (AD),^89,90^ and isomerization of tau at Asp387 in the peptide TDHGAEIVYK was recently found to be a stronger predictor of dementia in AD than aberrant protein aggregation.^26^ However, the rate of isomerization of tau is unknown, and the concentration of tau was not sufficient for detection in our cellular extracts. Therefore, to determine the rate of isomerization of tau, intact recombinant protein was aged *in vitro*. In addition, tau was incubated alone in buffer and in the presence heparin to induce aggregation.^72^

The progression of the isomerization for TDHGAEIVYK extracted from tryptic digests of incubated tau is shown in Figure 7a. Under standard PBS buffer conditions, tau should exist largely in the monomeric state with primarily disordered structure.^91^ These conditions lead to ∼27% isomerization by Day 50. To place this amount of isomerization in context with the incubation results described above, we plot the percent isomerization versus protein size (# of amino acids) in Fig. 7b. The results are additionally labeled by DG/nonDG status, where again these two groups clearly separate. Although there is no clear trend between protein size and isomerization, the datapoint for monomerized tau is significantly higher than any other nonDG peptide. In fact, extent of isomerization is ∼3x greater than the average for all observed non-DG sites and 1.35x larger than the next closest value. Tau is an intrinsically disordered protein, meaning it does not adopt a regular folded structure in isolation,^92^ which should provide the backbone flexibility needed for isomerization. The C-terminal residue is His, which is large and capable of steric interference but is also well-known for catalyzing acid/base reactions. We did observe isomerization at another DH site in the peptide YAVTTGDHGIIR from COPA, but the rate was significantly lower due to the local beta sheet structure, and isomerization only reached 7% by 50 days of aging. Therefore, the properties that enable rapid isomerization of monomeric tau are not immediately obvious, but the strong anti-correlation between isomerization and brain function is likely due to tau’s ability to isomerize rapidly and serve as a reporter for disrupted proteostasis.

**Figure 7.**
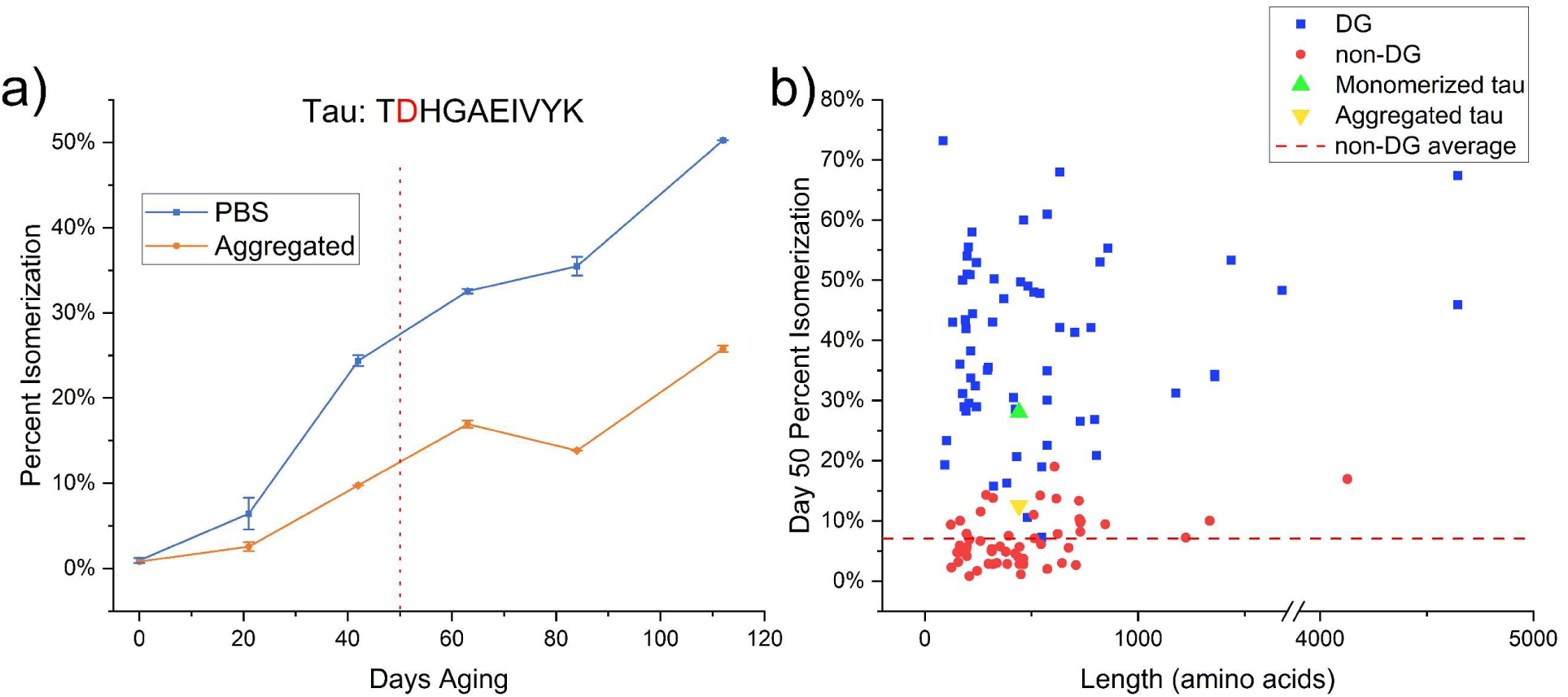
Progression of isomerization for the peptide TDHGAEIVYK from recombinant tau. (a) Protein aged only in PBS buffer is shown in blue, while protein in aggregating conditions is shown in orange. Red-dotted line corresponds to 50 days of aging. Error bars are derived from technical replicates. (b) percent isomerization compared to the length of the protein in amino acids, with DG isomers in blue, non-DG isomers in red, and the average %isomerization for non-DG isomers in the red, dashed line. Percent isomerization at Day 50 for TDHGAEIVYK from tau shown, with monomeric tau in green and aggregated tau in yellow.

To examine the potential effects of aggregation on tau isomerization, samples were incubated in PBS buffer with the addition of 10 µM heparin to induce aggregation.^72^ Although heparin-induced aggregation may not faithfully reproduce the specific aggregated states associated with AD or other tauopathies,^93^ any aggregation should restrict the conformational freedom of the protein backbone. Indeed, as seen in Fig. 7a, heparin-induced aggregation does slow tau isomerization, likely due to loss of conformational flexibility. Our data also suggest that heparin-induced aggregation may yield inconsistent amounts of aggregation or inconsistently aggregated structures. To clarify, the apparent isomerization at Day 84 is less than Day 64, which is not consistent with increased isomerization over time (yet replicate measurements for each day are highly reproducible and suggest the difference between the values is outside measurement error). However, each timepoint in Fig.7a was derived from a separate incubation, meaning that either more aggregation or more restrictive aggregates appear to have formed in the Day 84 sample. Since heparin incubation is a poor mimic for disease-relevant tau aggregation, our intent was primarily to establish whether aggregation could possibly slow isomerization, and the data in Fig. 7a clearly show that this is the case.

## Conclusions

Using aged cell lysates as an *in vitro* model of a complex isomer-containing proteome, DIA-based LC-MS allows for straightforward identification and quantification of Asp isomer sites in many proteins. Our results illustrate that multiple levels of protein structure fundamentally influence the rate of Asp isomerization *in vitro*. Crowding of the C-terminal amide decreases isomerization rate due to steric hindrance of succinimide ring formation. Increased backbone rigidity due to secondary structures, such as alpha helices and beta sheets, and protein complex formation also reduces isomerization. Asp isomerization was shown to not respond to base catalysis in intact proteins, a fundamental difference from the similar Asn deamidation. The determination of the principal factors underlying isomerization allows for prediction of which proteins are susceptible. Proteins associated with neurodegenerative diseases, such as alpha synuclein, huntingtin, and tau, contain a high density of Asp residues and are intrinsically disordered, so we would predict that they would likely isomerize at a very fast rate. As a case study, *in vitro* aging of recombinant tau also revealed that isomerization of sites with disease- dependence in Alzheimer’s disease proceeds at a faster than expected rate based on sequence likely owed to other structural properties of tau such as lack of crowding and a flexible backbone. Given that isomeric peptides are not degraded efficiently in the lysosome, the fast isomerization rate of tau provides a rationale for observed lysosomal storage in Alzheimer’s disease. Slower isomerization of tau in the aggregated condition suggests that isomerization may occur in soluble tau prior to aggregation, but analysis of insoluble tau in Alzheimer’s disease tissue and disease-specific aggregates is needed. Additionally, tau’s function of binding microtubules leaves questions regarding the conformational state in which isomerization proceeds *in vivo*. Overall, the detection of Asp isomers in the context of neurodegenerative disease has necessitated an understanding of the trends that underlie isomerization, and our results clearly illustrate that consideration of multiple levels of protein structure provides a framework to predict the propensity for Asp isomerization in a biological context.

## Supporting Information

Example of sequence isomer aging profile, comparison between the total number of peptides and number of isomers for each residue pair, structures of highly crowded low isomerization DG sites, structures of selected isomer sites, table of all quantifiably isomerized peptides, table of isomers impacted by aging in Native lysis buffers, table of isomers impacted by aging in ammonia-containing, higher pH lysis buffers.

## Acknowledgements

The authors gratefully acknowledge funding from the NIH (R01AG066626). V.O. was supported by a fellowship from the California Louis Stokes Alliance for Minority Participation (NSF#1826900)

## References

1 Cambridge, S. B.; Gnad, F.; Nguyen, C.; Bermejo, J. L.; Krüger, M.; Mann, M. Systems-Wide Proteomic Analysis in Mammalian Cells Reveals Conserved, Functional Protein Turnover. J. Proteome Res. 2011, 10 (12), 5275–5284.

2 Kluever, V.; Russo, B.; Mandad, S.; Kumar, N. H.; Alevra, M.; Ori, A.; Rizzoli, S. O.; Urlaub, H.; Schneider, A.; Fornasiero, E. F. Protein Lifetimes in Aged Brains Reveal a Proteostatic Adaptation Linking Physiological Aging to Neurodegeneration. Sci. Adv. 2022, 8 (20), eabn4437.

3 Price, J. C.; Guan, S.; Burlingame, A.; Prusiner, S. B.; Ghaemmaghami, S. Analysis of Proteome Dynamics in the Mouse Brain. Proc. Natl. Acad. Sci. U. S. A. 2010, 107 (32), 14508– 14513.

4 Truscott, R. J. W.; Schey, K. L.; Friedrich, M. G. Old Proteins in Man: A Field in Its Infancy. Trends Biochem. Sci. 2016, 41 (8), 654–664.

5 Savas, J. N.; Toyama, B. H.; Xu, T.; Yates, J. R.; Hetzer, M. W. Extremely Long-Lived Nuclear Pore Proteins in the Rat Brain. Science 2012, 335 (6071), 942.

6 Thayer, N. H.; Leverich, C. K.; Fitzgibbon, M. P.; Nelson, Z. W.; Henderson, K. A.; Gafken, P. R.; Hsu, J. J.; Gottschling, D. E. Identification of Long-Lived Proteins Retained in Cells Undergoing Repeated Asymmetric Divisions. Proc. Natl. Acad. Sci. U. S. A. 2014, 111 (39), 14019–14026.

7 Fornasiero, E. F.; Mandad, S.; Wildhagen, H.; Alevra, M.; Rammner, B.; Keihani, S.; Opazo, F.; Urban, I.; Ischebeck, T.; Sakib, M. S.; Fard, M. K.; Kirli, K.; Centeno, T. P.; Vidal, R. O.; Rahman, R.-U.; Benito, E.; Fischer, A.; Dennerlein, S.; Rehling, P.; Feussner, I.; Bonn, S.; Simons, M.; Urlaub, H.; Rizzoli, S. O. Precisely Measured Protein Lifetimes in the Mouse Brain Reveal Differences across Tissues and Subcellular Fractions. Nat. Commun. 2018, 9 (1), 4230.

8 Samuel Zigler, J.; Goosey, J. Aging of Protein Molecules: Lens Crystallins as a Model System. Trends Biochem. Sci. 1981, 6, 133–136.

9 Fujii, N.; Ishibashi, Y.; Satoh, K.; Fujino, M.; Harada, K. Simultaneous Racemization and Isomerization at Specific Aspartic Acid Residues in αB-Crystallin from the Aged Human Lens. *Biochim. Biophys. Acta*, Protein Struct. Mol. Enzymol. 1994, 1204 (2), 157–163.

10. Toyama, B. H.; Hetzer, M. W. Protein Homeostasis: Live Long, Won’t Prosper. Nat. Rev. Mol. Cell Biol. 2013, 14 (1), 55–61.

11. Cloos, P. A. C.; Christgau, S. Non-Enzymatic Covalent Modifications of Proteins: Mechanisms, Physiological Consequences and Clinical Applications. Matrix Biol. 2002, 21 (1), 39–52.

12 Truscott, R. J. W. Age-Related Nuclear Cataract—Oxidation Is the Key. Exp. Eye Res. 2005, 80 (5), 709–725.

13 Tonie Wright, H.; Urry, D. W. Nonenzymatic Deamidation of Asparaginyl and Glutaminyl Residues in Protein. Crit. Rev. Biochem. Mol. Biol. 1991, 26 (1), 1–52.

14 Höhn, A.; König, J.; Grune, T. Protein Oxidation in Aging and the Removal of Oxidized Proteins. J. Proteomics 2013, 92, 132–159.

15 Kim, Y. H.; Kapfer, D. M.; Boekhorst, J.; Lubsen, N. H.; Bächinger, H. P.; Shearer, T. R.; David, L. L.; Feix, J. B.; Lampi, K. J. Deamidation, but Not Truncation, Decreases the Urea Stability of a Lens Structural Protein, βB1-Crystallin. Biochemistry 2002, *41* (47), 14076–14084.

16 Lyon, Y. A.; Collier, M. P.; Riggs, D. L.; Degiacomi, M. T.; Benesch, J. L. P.; Julian, R. R. Structural and Functional Consequences of Age-Related Isomerization in α-Crystallins. J. Biol. Chem. 2019, 294 (19), 7546–7555.

17 Silzel, J. W.; Ben-Nissan, G.; Tang, J.; Sharon, M.; Julian, R. R. Influence of Asp Isomerization on Trypsin and Trypsin-like Proteolysis. Anal. Chem. 2022, 94 (44), 15288–15296.

18 Pandey, G.; Julian, R. R. LC–MS Reveals Isomeric Inhibition of Proteolysis by Lysosomal Cathepsins. Analysis Sensing 2022, 2 (4).

19 Geiger, T.; Clarke, S. Deamidation, Isomerization, and Racemization at Asparaginyl and Aspartyl Residues in Peptides. Succinimide-Linked Reactions That Contribute to Protein Degradation. J. Biol. Chem. 1987, 262 (2), 785–794.

20 Riggs, D. L.; Gomez, S. V.; Julian, R. R. Sequence and Solution Effects on the Prevalence of D-Isomers Produced by Deamidation. ACS Chem. Biol. 2017, 12 (11), 2875–2882.

21 Lambeth, T. R.; Riggs, D. L.; Talbert, L. E.; Tang, J.; Coburn, E.; Kang, A. S.; Noll, J.; Augello, C.; Ford, B. D.; Julian, R. R. Spontaneous Isomerization of Long-Lived Proteins Provides a Molecular Mechanism for the Lysosomal Failure Observed in Alzheimer’s Disease. ACS Cent. Sci. 2019, 5 (8), 1387–1395.

22. Julian, R. R. How Isomerization and Epimerization in Long-Lived Proteins Affect Lysosomal Degradation and Proteostasis. In Long-lived Proteins in Human Aging and Disease; John Wiley & Sons, Ltd, 2021; pp 175–188.

23 Van Acker, Z. P.; Bretou, M.; Annaert, W. Endo-Lysosomal Dysregulations and Late-Onset Alzheimer’s Disease: Impact of Genetic Risk Factors. Mol. Neurodegener. 2019, 14 (1), 20.

24 Nixon, R. A. The Aging Lysosome: An Essential Catalyst for Late-Onset Neurodegenerative Diseases. *Biochim. Biophys. Acta*, Proteins Proteomics 2020, 1868 (9), 140443.

25 Fleming, A.; Bourdenx, M.; Fujimaki, M.; Karabiyik, C.; Krause, G. J.; Lopez, A.; Martín-Segura, A.; Puri, C.; Scrivo, A.; Skidmore, J.; Son, S. M.; Stamatakou, E.; Wrobel, L.; Zhu, Y.; Cuervo, A. M.; Rubinsztein, D. C. The Different Autophagy Degradation Pathways and Neurodegeneration. Neuron 2022, 110 (6), 935–966.

26 Hubbard, E. E.; Heil, L. R.; Merrihew, G. E.; Chhatwal, J. P.; Farlow, M. R.; McLean, C. A.; Ghetti, B.; Newell, K. L.; Frosch, M. P.; Bateman, R. J.; Larson, E. B.; Keene, C. D.; Perrin, R. J.; Montine, T. J.; MacCoss, M. J.; Julian, R. R. Does Data-Independent Acquisition Data Contain Hidden Gems? A Case Study Related to Alzheimer’s Disease. J. Proteome Res. 2022, 21 (1), 118–131.

27 Yang, H.; Fung, E. Y. M.; Zubarev, A. R.; Zubarev, R. A. Toward Proteome-Scale Identification and Quantification of Isoaspartyl Residues in Biological Samples. J. Proteome Res. 2009, 8 (10), 4615–4621.

28 Sargaeva, N. P.; Lin, C.; O’Connor, P. B. Unusual Fragmentation of β-Linked Peptides by ExD Tandem Mass Spectrometry. J. Am. Soc. Mass Spectrom. 2011, 22 (3), 480–491.

29 Ni, W.; Dai, S.; Karger, B. L.; Zhou, Z. S. Analysis of Isoaspartic Acid by Selective Proteolysis with Asp-N and Electron Transfer Dissociation Mass Spectrometry. Anal. Chem. 2010, 82 (17), 7485–7491.

30 Tao, Y.; Quebbemann, N. R.; Julian, R. R. Discriminating D-Amino Acid-Containing Peptide Epimers by Radical-Directed Dissociation Mass Spectrometry. Anal. Chem. 2012, 84 (15), 6814–6820.

31 Wu, H.-T.; Riggs, D. L.; Lyon, Y. A.; Julian, R. R. Statistical Framework for Identifying Differences in Similar Mass Spectra: Expanding Possibilities for Isomer Identification. Anal. Chem. 2023, 95 (17), 6996–7005.

32 M. Edwards, H.; Wu, H.-T. R. Julian, R.; P. Jackson, G. Differentiating Aspartic Acid Isomers and Epimers with Charge Transfer Dissociation Mass Spectrometry (CTD-MS). Analyst 2022, 147 (6), 1159–1168.

33 Wu, H.-T.; R. Julian, R. Two-Dimensional Identification and Localization of Isomers in Crystallin Peptides Using TWIM-MS. Analyst 2020, 145 (15), 5232–5241.

34 Jia, C.; Lietz, C. B.; Yu, Q.; Li, L. Site-Specific Characterization of d-Amino Acid Containing Peptide Epimers by Ion Mobility Spectrometry. Anal. Chem. 2014, 86 (6), 2972–2981.

35 Zheng, X.; Deng, L.; Baker, E. S.; Ibrahim, Y. M.; Petyuk, V. A.; Smith, R. D. Distinguishing D- and L-Aspartic and Isoaspartic Acids in Amyloid β Peptides with Ultrahigh Resolution Ion Mobility Spectrometry. Chem. Commun. (Cambridge, U. K.) 2017, 53 (56), 7913–7916.

36 Silzel, J. W.; Lambeth, T. R.; Julian, R. R. PIMT-Mediated Labeling of l-Isoaspartic Acid with Tris Facilitates Identification of Isomerization Sites in Long-Lived Proteins. J. Am. Soc. Mass Spectrom. 2022, 33 (3), 548–556.

37 Lyon, Y. A.; Sabbah, G. M.; Julian, R. R. Differences in α-Crystallin Isomerization Reveal the Activity of Protein Isoaspartyl Methyltransferase (PIMT) in the Nucleus and Cortex of Human Lenses. Exp. Eye Res. 2018, 171, 131–141.

38 Lyon, Y. A.; Sabbah, G. M.; Julian, R. R. Identification of Sequence Similarities among Isomerization Hotspots in Crystallin Proteins. J. Proteome Res. 2017, 16 (4), 1797–1805.

39 Bischoff, R.; Kolbe, H. V. J. Deamidation of Asparagine and Glutamine Residues in Proteins and Peptides: Structural Determinants and Analytical Methodology. J. Chromatogr. B: Biomed. Sci. Appl. 1994, 662 (2), 261–278.

40 Yang, H.; Zubarev, R. A. Mass Spectrometric Analysis of Asparagine Deamidation and Aspartate Isomerization in Polypeptides. Electrophoresis 2010, 31 (11), 1764–1772.

41 Robinson, N. E.; Robinson, A. B. Molecular Clocks. Proc. Natl. Acad. Sci. U. S. A. 2001, 98 (3), 944–949.

42 Robinson, N. e.; Robinson, Z. w.; Robinson, B. r.; Robinson, A. l.; Robinson, J. a.; Robinson, M. l.; Robinson, A. b. Structure-Dependent Nonenzymatic Deamidation of Glutaminyl and Asparaginyl Pentapeptides. J. Pept. Res. 2004, 63 (5), 426–436.

43 Robinson, N. E.; Robinson, A. B. Prediction of Protein Deamidation Rates from Primary and Three-Dimensional Structure. Proc. Natl. Acad. Sci. U. S. A. 2001, 98 (8), 4367–4372.

44 Robinson, N. E.; Robinson, A. B. Deamidation of Human Proteins. Proc. Natl. Acad. Sci. U. S. A. 2001, 98 (22), 12409–12413.

45 Robinson, N. E.; Robinson, M. L.; Schulze, S. E. S.; Lai, B. T.; Gray, H. B. Deamidation of α-Synuclein. Protein Sci. 2009, 18 (8), 1766–1773.

46 Kuge, K.; Brack, A.; Fujii, N. Conformation-Dependent Racemization of Aspartyl Residues in Peptides. Chem. - Eur. J. 2007, 13 (19), 5617–5621.

47 Zhang, W.; Czupryn, M. J. Analysis of Isoaspartate in a Recombinant Monoclonal Antibody and Its Charge Isoforms. J. Pharm. Biomed. Anal. 2003, 30 (5), 1479–1490.

48 Yokoyama, H.; Mizutani, R.; Noguchi, S.; Hayashida, N. Structural and Biochemical Basis of the Formation of Isoaspartate in the Complementarity-Determining Region of Antibody 64M-5 Fab. Sci. Rep. 2019, 9 (1), 18494.

49 Wakankar, A. A.; Borchardt, R. T.; Eigenbrot, C.; Shia, S.; Wang, Y. J.; Shire, S. J.; Liu, J. L. Aspartate Isomerization in the Complementarity-Determining Regions of Two Closely Related Monoclonal Antibodies. Biochemistry 2007, 46 (6), 1534–1544.

50 Dick, L. W.; Qiu, D.; Cheng, K.-C. Identification and Measurement of Isoaspartic Acid Formation in the Complementarity Determining Region of a Fully Human Monoclonal Antibody. J. Chromatogr. B 2009, 877 (30), 3841–3849.

51 Sreedhara, A.; Cordoba, A.; Zhu, Q.; Kwong, J.; Liu, J. Characterization of the Isomerization Products of Aspartate Residues at Two Different Sites in a Monoclonal Antibody. Pharm. Res. 2012, 29 (1), 187–197.

52 Yi, M.; Sun, J.; Sun, H.; Wang, Y.; Hou, S.; Jiang, B.; Xie, Y.; Ji, R.; Xue, L.; Ding, X.; Song, X.; Xu, A.; Huang, C.; Quan, Q.; Song, J. Identification and Characterization of an Unexpected Isomerization Motif in CDRH2 That Affects Antibody Activity. mAbs 2023, 15 (1), 2215364.

53 Tao, Y.; Julian, R. R. Identification of Amino Acid Epimerization and Isomerization in Crystallin Proteins by Tandem LC-MS. Anal. Chem. 2014, 86 (19), 9733–9741.

54 Bloemendal, H.; de Jong, W.; Jaenicke, R.; Lubsen, N. H.; Slingsby, C.; Tardieu, A. Ageing and Vision: Structure, Stability and Function of Lens Crystallins. Prog. Biophys. Mol. Biol. 2004, 86 (3), 407–485.

55 Roskamp, K. W.; Paulson, C. N.; Brubaker, W. D.; Martin, R. W. Function and Aggregation in Structural Eye Lens Crystallins. Acc. Chem. Res. 2020, 53 (4), 863–874.

56 Grosas, A. B.; Carver, J. A. Eye Lens Crystallins: Remarkable Long-Lived Proteins. In Long-lived Proteins in Human Aging and Disease; John Wiley & Sons, Ltd, 2021; pp 59–96.

57 Zhu, X.; Korlimbinis, A.; Truscott, R. J. W. Age-Dependent Denaturation of Enzymes in the Human Lens: A Paradigm for Organismic Aging? Rejuvenation Res. 2010, 13 (5), 553–560.

58 Egertson, J. D.; MacLean, B.; Johnson, R.; Xuan, Y.; MacCoss, M. J. Multiplexed Peptide Analysis Using Data-Independent Acquisition and Skyline. Nat. Protoc. 2015, 10 (6), 887–903.

59 Amodei, D.; Egertson, J.; MacLean, B. X.; Johnson, R.; Merrihew, G. E.; Keller, A.; Marsh, D.; Vitek, O.; Mallick, P.; MacCoss, M. J. Improving Precursor Selectivity in Data-Independent Acquisition Using Overlapping Windows. J. Am. Soc. Mass Spectrom. 2019, 30 (4), 669–684.

60 Egertson, J. D.; Kuehn, A.; Merrihew, G. E.; Bateman, N. W.; MacLean, B. X.; Ting, Y. S.; Canterbury, J. D.; Marsh, D. M.; Kellmann, M.; Zabrouskov, V.; Wu, C. C.; MacCoss, M. J. Multiplexed MS/MS for Improved Data-Independent Acquisition. Nat. Methods 2013, 10 (8), 744–746.

61 Adusumilli, R.; Mallick, P. Data Conversion with ProteoWizard msConvert. In Proteomics: Methods and Protocols; Comai, L., Katz, J. E., Mallick, P., Eds.; Springer: New York, NY, 2017; pp 339–368.

62 Chambers, M. C.; Maclean, B.; Burke, R.; Amodei, D.; Ruderman, D. L.; Neumann, S.; Gatto, L.; Fischer, B.; Pratt, B.; Egertson, J.; Hoff, K.; Kessner, D.; Tasman, N.; Shulman, N.; Frewen, B.; Baker, T. A.; Brusniak, M.-Y.; Paulse, C.; Creasy, D.; Flashner, L.; Kani, K.; Moulding, C.; Seymour, S. L.; Nuwaysir, L. M.; Lefebvre, B.; Kuhlmann, F.; Roark, J.; Rainer, P.; Detlev, S.; Hemenway, T.; Huhmer, A.; Langridge, J.; Connolly, B.; Chadick, T.; Holly, K.; Eckels, J.; Deutsch, E. W.; Moritz, R. L.; Katz, J. E.; Agus, D. B.; MacCoss, M.; Tabb, D. L.; Mallick, P. A Cross-Platform Toolkit for Mass Spectrometry and Proteomics. Nat. Biotechnol. 2012, 30 (10), 918–920.

63 Searle, B. C.; Pino, L. K.; Egertson, J. D.; Ting, Y. S.; Lawrence, R. T.; MacLean, B. X.; Villén, J.; MacCoss, M. J. Chromatogram Libraries Improve Peptide Detection and Quantification by Data Independent Acquisition Mass Spectrometry. Nat. Commun. 2018, 9 (1), 5128.

64 Ting, Y. S.; Egertson, J. D.; Bollinger, J. G.; Searle, B. C.; Payne, S. H.; Noble, W. S.; MacCoss, M. J. PECAN: Library-Free Peptide Detection for Data-Independent Acquisition Tandem Mass Spectrometry Data. Nat. Methods 2017, 14 (9), 903–908.

65 Pino, L. K.; Just, S. C.; MacCoss, M. J.; Searle, B. C. Acquiring and Analyzing Data Independent Acquisition Proteomics Experiments without Spectrum Libraries. Mol. Cell. Proteomics 2020, 19 (7), 1088–1103.

66 Gessulat, S.; Schmidt, T.; Zolg, D. P.; Samaras, P.; Schnatbaum, K.; Zerweck, J.; Knaute, T.; Rechenberger, J.; Delanghe, B.; Huhmer, A.; Reimer, U.; Ehrlich, H.-C.; Aiche, S.; Kuster, B.; Wilhelm, M. Prosit: Proteome-Wide Prediction of Peptide Tandem Mass Spectra by Deep Learning. Nat. Methods 2019, 16 (6), 509–518.

67 Käll, L.; Canterbury, J. D.; Weston, J.; Noble, W. S.; MacCoss, M. J. Semi-Supervised Learning for Peptide Identification from Shotgun Proteomics Datasets. Nat. Methods 2007, 4 (11), 923–925.

68 Käll, L.; Storey, J. D.; MacCoss, M. J.; Noble, W. S. Posterior Error Probabilities and False Discovery Rates: Two Sides of the Same Coin. J. Proteome Res. 2008, 7 (1), 40–44.

69 MacLean, B.; Tomazela, D. M.; Shulman, N.; Chambers, M.; Finney, G. L.; Frewen, B.; Kern, R.; Tabb, D. L.; Liebler, D. C.; MacCoss, M. J. Skyline: An Open Source Document Editor for Creating and Analyzing Targeted Proteomics Experiments. Bioinformatics 2010, *26* (7), 966–968.

70 Pino, L. K.; Searle, B. C.; Bollinger, J. G.; Nunn, B.; MacLean, B.; MacCoss, M. J. The Skyline Ecosystem: Informatics for Quantitative Mass Spectrometry Proteomics. Mass Spectrom. Rev. 2020, 39 (3), 229–244.

71 Qi, J.; Xiao, Y. Prediction of Nonregular Secondary Structures of Proteins Based on Wavelet Analysis. Chin. Sci. Bull. 2002, 47 (11), 918–923.

72 Montgomery, K. M.; Carroll, E. C.; Thwin, A. C.; Quddus, A. Y.; Hodges, P.; Southworth, D. R.; Gestwicki, J. E. Chemical Features of Polyanions Modulate Tau Aggregation and Conformational States. J. Am. Chem. Soc. 2023, 145 (7), 3926–3936.

73 Ngoka, L. C. Sample Prep for Proteomics of Breast Cancer: Proteomics and Gene Ontology Reveal Dramatic Differences in Protein Solubilization Preferences of Radioimmunoprecipitation Assay and Urea Lysis Buffers. Proteome Sci. 2008, 6 (1), 30.

74 Qi, J.; Xiao, Y. Prediction of Nonregular Secondary Structures of Proteins Based on Wavelet Analysis. Chin. Sci. Bull. 2002, 47 (11), 918–923.

75 Ippolito, J. A.; Alexander, R. S.; Christianson, D. W. Hydrogen Bond Stereochemistry in Protein Structure and Function. J. Mol. Biol. 1990, 215 (3), 457–471.

76 Kemp, M. T.; Lewandowski, E. M.; Chen, Y. Low Barrier Hydrogen Bonds in Protein Structure and Function. *Biochim. Biophys. Acta*, Proteins Proteomics 2021, 1869 (1), 140557.

77 Kortemme, T.; Morozov, A. V.; Baker, D. An Orientation-Dependent Hydrogen Bonding Potential Improves Prediction of Specificity and Structure for Proteins and Protein–Protein Complexes. J. Mol. Biol. 2003, 326 (4), 1239–1259.

78 Pace, C. N.; Fu, H.; Lee Fryar, K.; Landua, J.; Trevino, S. R.; Schell, D.; Thurlkill, R. L.; Imura, S.; Scholtz, J. M.; Gajiwala, K.; Sevcik, J.; Urbanikova, L.; Myers, J. K.; Takano, K.; Hebert, E. J.; Shirley, B. A.; Grimsley, G. R. Contribution of Hydrogen Bonds to Protein Stability. Protein Sci. 2014, 23 (5), 652–661.

79 Kosky, A. A.; Razzaq, U. O.; Treuheit, M. J.; Brems, D. N. The Effects of Alpha-Helix on the Stability of Asn Residues: Deamidation Rates in Peptides of Varying Helicity. Protein Sci. 1999, 8 (11), 2519–2523.

80 Vijayakumar, S.; Vishveshwara, S.; Ravishanker, G.; Beveridge, D. L. Differential Stability of Beta-Sheets and Alpha-Helices in Beta-Lactamase: A High Temperature Molecular Dynamics Study of Unfolding Intermediates. Biophys. J. 1993, 65 (6), 2304–2312.

81 Jones, S.; Thornton, J. M. Principles of Protein-Protein Interactions. Proc. Natl. Acad. Sci. U. S. A. 1996, 93 (1), 13–20.

82 Keskin, O.; Gursoy, A.; Ma, B.; Nussinov, R. Principles of Protein−Protein Interactions: What Are the Preferred Ways For Proteins To Interact? Chem. Rev. 2008, 108 (4), 1225–1244.

83 Mei, L.; Montoya, M. R.; Quanrud, G. M.; Tran, M.; Villa-Sharma, A.; Huang, M.; Genereux, J. C. Bait Correlation Improves Interactor Identification by Tandem Mass Tag-Affinity Purification-Mass Spectrometry. J. Proteome Res. 2020, 19 (4), 1565–1573.

84 Schröder, E.; Littlechil*, J. A.; Lebedev, A. A.; Errington, N.; Vagin, A. A.; Isupov, M. N. Crystal Structure of Decameric 2-Cys Peroxiredoxin from Human Erythrocytes at 1.7Å Resolution. Structure 2000, 8 (6), 605–615.

85 Anger, A. M.; Armache, J.-P.; Berninghausen, O.; Habeck, M.; Subklewe, M.; Wilson, D. N.; Beckmann, R. Structures of the Human and Drosophila 80S Ribosome. Nature 2013, 497 (7447), 80–85.

86 Riggs, D. L.; Gomez, S. V.; Julian, R. R. Sequence and Solution Effects on the Prevalence of D-Isomers Produced by Deamidation. ACS Chem. Biol. 2017, 12 (11), 2875–2882.

87 Tyler-Cross, R.; Schirch, V. Effects of Amino Acid Sequence, Buffers, and Ionic Strength on the Rate and Mechanism of Deamidation of Asparagine Residues in Small Peptides. J. Biol. Chem. 1991, 266 (33), 22549–22556.

88 Oliyai, C.; Borchardt, R. T. Chemical Pathways of Peptide Degradation. IV. Pathways, Kinetics, and Mechanism of Degradation of an Aspartyl Residue in a Model Hexapeptide. Pharm. Res. 1993, 10 (1), 95–102.

89 Mamun, A. A.; Uddin, M. S.; Mathew, B.; Ashraf, G. M. Toxic Tau: Structural Origins of Tau Aggregation in Alzheimer’s Disease. Neural Regener. Res. 2020, 15 (8), 1417.

90 Lasagna-Reeves, C. A.; Castillo-Carranza, D. L.; Sengupta, U.; Sarmiento, J.; Troncoso, J.; Jackson, G. R.; Kayed, R. Identification of Oligomers at Early Stages of Tau Aggregation in Alzheimer’s Disease. FASEB J. 2012, 26 (5), 1946–1959.

91 Iqbal, K.; Liu, F.; Gong, C.-X.; Grundke-Iqbal, I. Tau in Alzheimer Disease and Related Tauopathies. Curr. Alzheimer Res. 2010, 7 (8), 656–664.

92 Kadavath, H.; Hofele, R. V.; Biernat, J.; Kumar, S.; Tepper, K.; Urlaub, H.; Mandelkow, E.; Zweckstetter, M. Tau Stabilizes Microtubules by Binding at the Interface between Tubulin Heterodimers. Proc. Natl. Acad. Sci. U. S. A. 2015, 112 (24), 7501–7506.

93 Zhang, W.; Falcon, B.; Murzin, A. G.; Fan, J.; Crowther, R. A.; Goedert, M.; Scheres, S. H. Heparin-Induced Tau Filaments Are Polymorphic and Differ from Those in Alzheimer’s and Pick’s Diseases. eLife 2019, 8, e43584.

