## Supporting Information for "Unusually Rapid Isomerization of Aspartic Acid in Tau"

Figure S1: (a) Example of Day 0 chromatogram and (b) %isomerization for sequence isomer across time points.

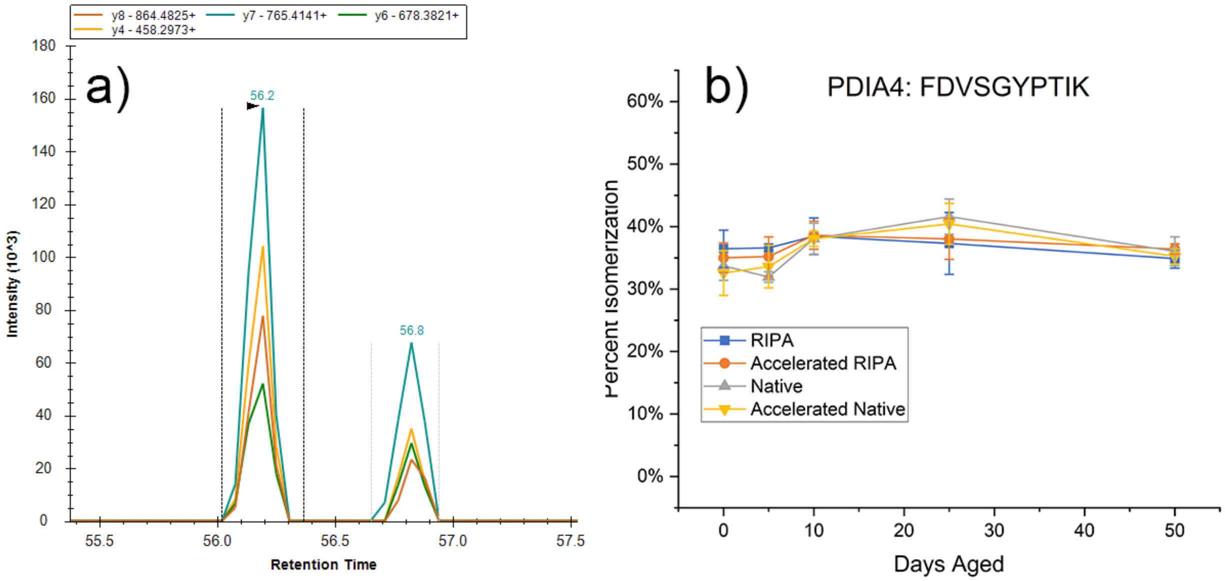

Figure S2: Comparison between the total number of peptides (orange) to the number of isomers (green) for each residue pair

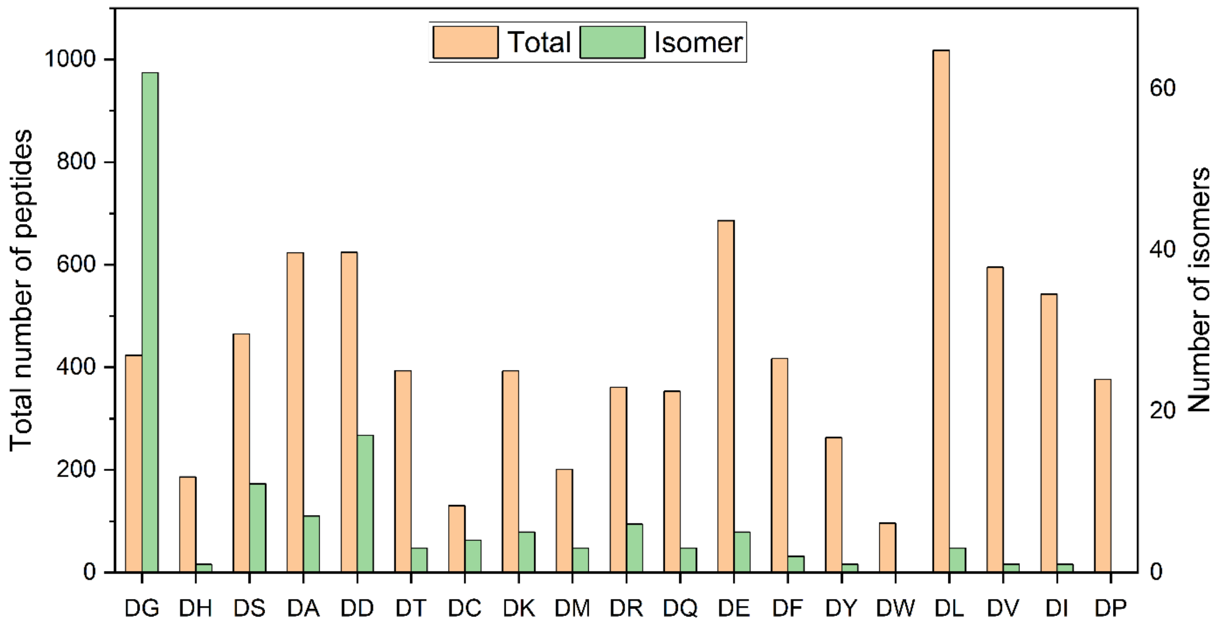

Figure S3: Examples of high crowding due to (a) short-range interactions or (b) long-range interactions that result in low isomerization in DPYL2

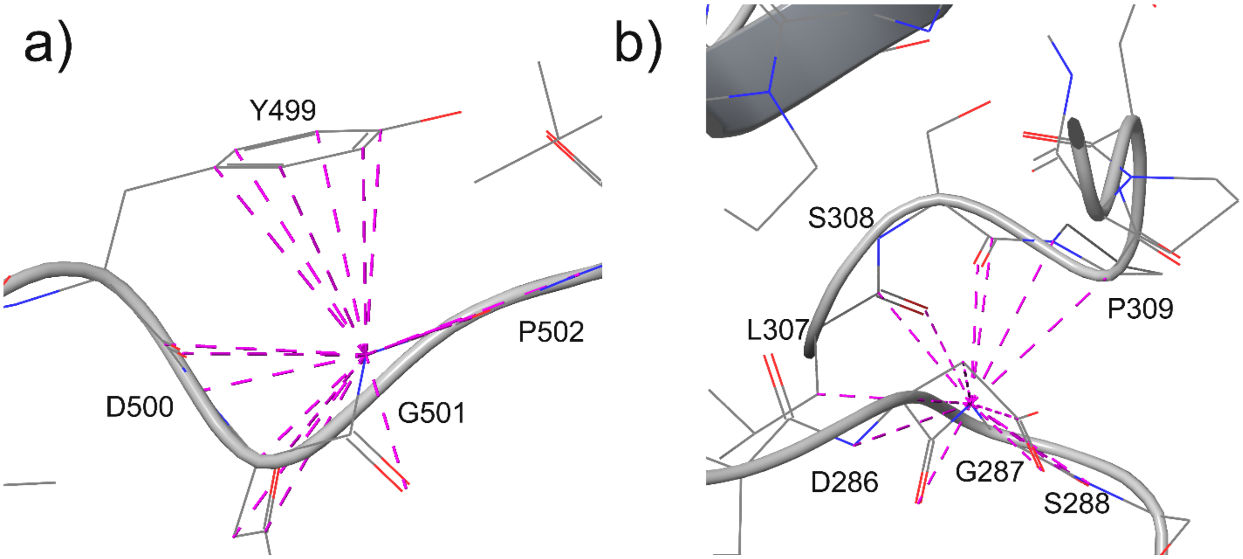

Figure S4: Structures of selected isomer sites that showed significant differences between RIPA and Native buffers (a-b) RAN: LVLVGDGGTGK (c-d) PRDX2: ATAVVDGAFK (e-f) HS90B: YHTSQSGDEMTSLSEYVSR (g-h) RS12: TALIHDGLAR

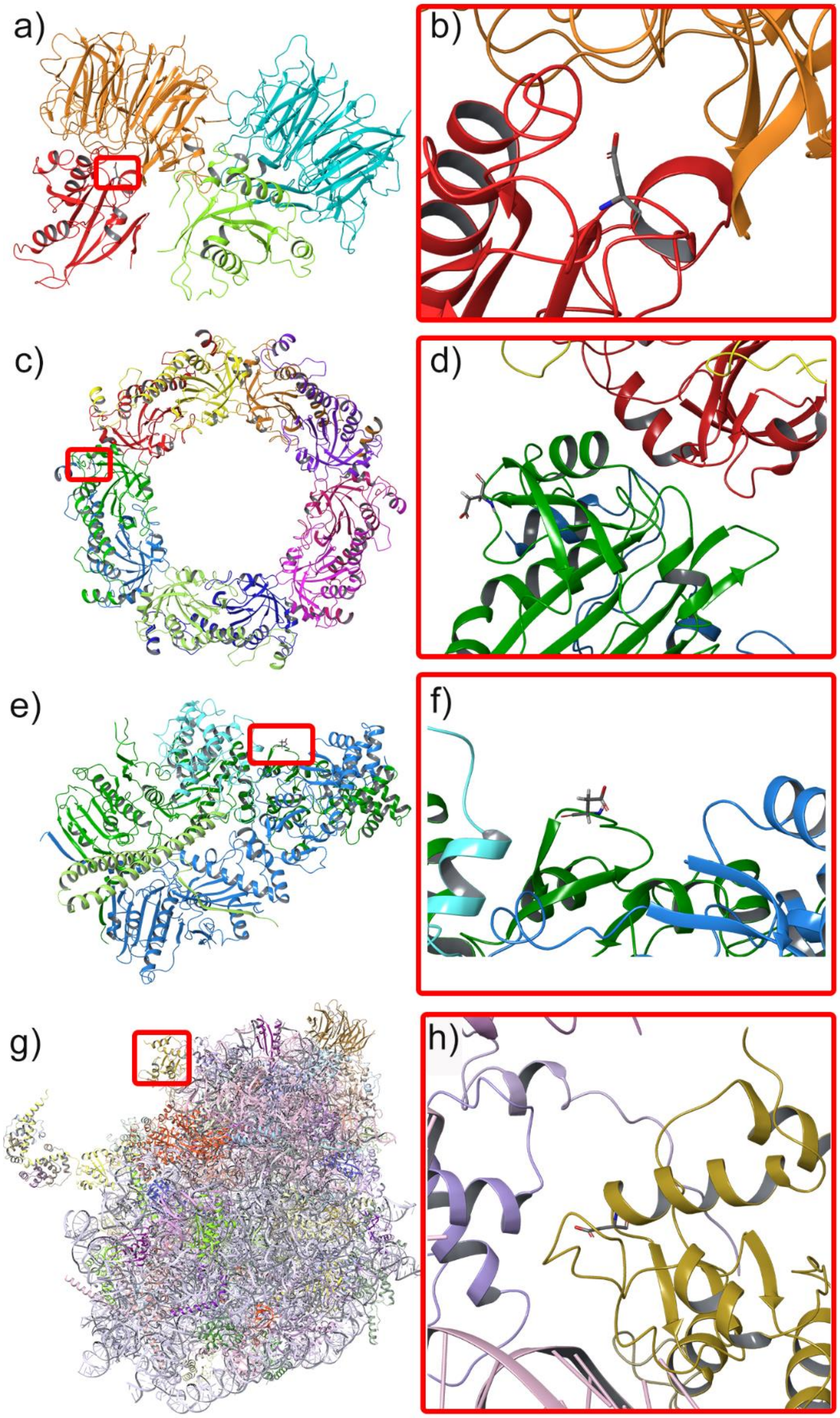

Table S1: List of identified Asp isomers, including protein, peptide sequence, and PDB accession number. AlphaFold IDs are listed when X-ray, EM, or NMR structures were unavailable containing the isomer site, written as “AF-uniprotID”.

| Protein | Peptide | PDB |
|---|---|---|
| sp\|P06576\|ATPB_HUMAN | TIAMDGTEGLVR | 8H9E |
| sp\|P36873\|PP1G_HUMAN | AHQVVEDGYEFFAK | 1IT6 |
| sp\|F5H284\|PAL4D_HUMAN | TEWLDGK | AF-F5H284 |
| sp\|P32322\|P5CR1_HUMAN | EGATVYATGTHAQVEDGR | 2GER |
| sp\|Q06830\|PRDX1_HUMAN | ATAVMPDGQFK | 2RII |
| sp\|Q06830\|PRDX1_HUMAN | LVQAFQFTDK | 2RII |
| sp\|Q9Y536\|PAL4A_HUMAN | IIPGFMCQGGDFTR | AF-Q9Y536 |
| sp\|P42766\|RL35_HUMAN | YKPLDLRPK | 4UG0 |
| sp\|P62826\|RAN_HUMAN | LVLVGDGGTGK | 1I2M |
| sp\|P15153\|RAC2_HUMAN | CVVVGDGAVGK | 1DS6 |
| sp\|P62913\|RL11_HUMAN | YDGIILPGK | 4UG0 |
| sp\|P22234\|PUR6_HUMAN | TKEVYELLDSPGK | 2H31 |
| sp\|P22234\|PUR6_HUMAN | IKAEYEGDGIPTVFVAVAGR | 2H31 |
| sp\|P23284\|PPIB_HUMAN | TAWLDGK | 1CYN |
| sp\|P50914\|RL14_HUMAN | ALVDGPCTQVR | 4UG0 |
| sp\|P25398\|RS12_HUMAN | TALIHDGLAR | 4V6X |
| sp\|P27635\|RL10_HUMAN | LIPDGCGVK | 5AJ0 |
| sp\|P84098\|RL19_HUMAN | LLADQAEAR | 4UG0 |
| sp\|P07437\|TBB5_HUMAN | ALTVPELTQQVFDAK | 5N5N |
| sp\|P49368\|TCPG_HUMAN | IVLLDSSLEYK | 6NR8 |
| sp\|P48643\|TCPE_HUMAN | IADGYEQAAR | 7NVL |
| sp\|P68363\|TBA1B_HUMAN | EDMAALEK | 5IJ0 |
| sp\|P04350\|TBB4A_HUMAN | ALTVPELTQQMFDAK | AF-P04350 |
| sp\|P62306\|RUXF_HUMAN | GVEEEEEDGEMRE | AF-P62306 |
| sp\|Q9Y230\|RUVB2_HUMAN | FVQCPDGELQK | AF-Q9Y230 |
| sp\|P23526\|SAHH_HUMAN | ESLIDGIKR | 1A7A |
| sp\|P55072\|TERA_HUMAN | IVSQLLTLMDGLK | 5C18 |
| sp\|P50990\|TCPQ_HUMAN | HEKEDGAISTIVLR | 7NVL |
| sp\|P22626\|ROA2_HUMAN | LTDCVVMR | 5EN1 |
| sp\|P32969\|RL9_HUMAN | FLDGIYVSEK | 4UG0 |
| sp\|P46777\|RL5_HUMAN | GAVDGGLSIPHSTK | 4UG0 |
| sp\|P62241\|RS8_HUMAN | ADGYVLEGK | 4UG0 |
| sp\|P46781\|RS9_HUMAN | IGVLDEGK | 4V6X |
| sp\|A6NKG5\|RTL1_HUMAN | LLHIHSADGQLHLLSR | AF-A6NKG5 |
| sp\|P23396\|RS3_HUMAN | FVADGIFK | 4UG0 |
| sp\|P23396\|RS3_HUMAN | ELAEDGYSGVEVR | 4UG0 |
| sp\|Q9BYD1\|RM13_HUMAN | IWYLLDGK | 3J7Y |

| | | |
|---|---|---|
| sp\|P61956\|SUMO2_HUMAN | VAGQDGSVVQFK | 1WM2 |
| sp\|P12956\|XRCC6_HUMAN | ILELDQFK | 1JEQ |
| sp\|P09936\|UCHL1_HUMAN | LGFEDGSVLK | 2ETL |
| sp\|P45880\|VDAC2_HUMAN | LTLSALVDGK | AF-P45880 |
| sp\|Q92841\|DDX17_HUMAN | GDGPICLVLAPTR | 6UV0 |
| sp\|P25705\|ATPA_HUMAN | VLSIGDGIAR | AF-P25705 |
| sp\|P25705\|ATPA_HUMAN | AVDSLVPIGR | AF-P25705 |
| sp\|Q16555\|DPYL2_HUMAN | GTVVYGEPITASLGTDGSHYWSK | 5X1A |
| sp\|Q16555\|DPYL2_HUMAN | GLYDGPVCEVSVTPK | 5X1A |
| sp\|P24534\|EF1B_HUMAN | SIQADGLVWGSSK | 1B64 |
| sp\|P17066\|HSP76_HUMAN | TTPSYVAFTDTER | 3FE1 |
| sp\|Q13347\|EIF3I_HUMAN | SYSSGGEDGYVR | 6ZON |
| sp\|Q14194\|DPYL1_HUMAN | GMYDGPVYEVPATPK | AF-Q14194 |
| sp\|P07900\|HS90A_HUMAN | YYTSASGDEMVSLK | 7RY0 |
| sp\|P52597\|HNRPF_HUMAN | VHIEIGPDGR | 2HGN |
| sp\|P61978\|HNRPK_HUMAN | LFQECCPHSTDR | AF-P61978 |
| sp\|P61978\|HNRPK_HUMAN | VVLIGGKPDR | AF-P61978 |
| sp\|P08238\|HS90B_HUMAN | YHTSQSGDEMTSLSEYVSR | 5FWK |
| sp\|P62269\|RS18_HUMAN | AGELTEDEVER | 4UG0 |
| sp\|P62937\|PPIA_HUMAN | VSFELFADKVPK | 1AK4 |
| sp\|P62937\|PPIA_HUMAN | KITIADCGQLE | 1AK4 |
| sp\|P32119\|PRDX2_HUMAN | ATAVVDGAFK | 1QMV |
| sp\|P38606\|VATA_HUMAN | LAEMPADSGYPAYLGAR | 6WLZ |
| sp\|P28331\|NDUS1_HUMAN | VLFLLGADGGCITR | 5XTB |
| sp\|P40926\|MDHM_HUMAN | GYLGPEQLPDCLK | 2DFD |
| sp\|P43243\|MATR3_HUMAN | GIDLLKK | AF-43243 |
| sp\|P26583\|HMGB2_HUMAN | SEHPGLSIGDTAK | AF-P26583 |
| sp\|P62310\|LSM3_HUMAN | GDGVVLVAPPLR | 6AH0 |
| sp\|P30613\|KPYR_HUMAN | GDLGIEIPAEK | 2VGB |
| sp\|P55795\|HNRH2_HUMAN | YGDGGSSFQSTTGHCVHMR | AF-P55795 |
| sp\|P38646\|GRP75_HUMAN | TTPSVVAFTADGER | 4KBO |
| sp\|P38646\|GRP75_HUMAN | SQVFSTAADGQTQVEIK | 3N8E |
| sp\|Q99497\|PARK7_HUMAN | DGLILTSR | 1J42 |
| sp\|O95433\|AHSA1_HUMAN | VFTTQELVQAFTHAPATLEADR | 7DME |
| sp\|P05141\|ADT2_HUMAN | GLGDCLVK | AF-P05141 |
| sp\|P63104\|1433Z_HUMAN | DSTLIMQLLR | 1QJA |
| sp\|P10809\|CH60_HUMAN | LSDGVAVLK | 6HT7 |
| sp\|Q9UJS0\|CMC2_HUMAN | ASGDSARPVLLQVAESAYR | AF-Q9UJS0 |
| sp\|Q92747\|ARC1A_HUMAN | DGVWKPTLVILR | AF-Q92747 |
| sp\|Q00610\|CLH1_HUMAN | LASTLVHLGEYQAAVDGAR | AF-Q00610 |
| sp\|O00154\|BACH_HUMAN | LMDEVAGIVAAR | 2QQ2 |

| | | |
|---|---|---|
| sp\|Q32P51\|RA1L2_HUMAN | EDSQRPGAHLTVK | AF-Q32P51 |
| sp\|Q32P51\|RA1L2_HUMAN | IEVIEIMTDR | AF-Q32P51 |
| sp\|Q9UQ80\|PA2G4_HUMAN | FDAMPFTLR | 2Q8K |
| sp\|Q9Y570\|PPME1_HUMAN | YWDGWFR | 3C5V |
| sp\|P17980\|PRS6A_HUMAN | QTYFLPVIGLVDAEK | 5GJQ |
| sp\|P78527\|PRKDC_HUMAN | SLGPPQGEEDSVPR | 5LUQ |
| sp\|P30101\|PDIA3_HUMAN | TADGIVSHLK | 7QNG |
| sp\|P12004\|PCNA_HUMAN | SEGFDTYR | 1AXC |
| sp\|P08237\|PFKAM_HUMAN | VLVVHDGFEGLAK | 4OMT |
| sp\|P19338\|NUCL_HUMAN | TEADAEKTFEEK | 2KRR |
| sp\|Q02878\|RL6_HUMAN | AVDSQILPK | 4UG0 |
| sp\|Q9H4L4\|SENP3_HUMAN | LGLLGALMAEDGVR | AF-Q9H4L4 |
| sp\|P62701\|RS4X_HUMAN | GIPHLVTHDAR | 4UG0 |
| sp\|Q7KZ85\|SPT6H_HUMAN | IMKIDIEK | 7OOP |
| sp\|Q9P2J5\|SYLC_HUMAN | GFYEGIMLVDGFK | 6KID |
| sp\|P45974\|UBP5_HUMAN | KQEVQAWDGEVR | AF-P45974 |
| sp\|Q16527\|CSRP2_HUMAN | TVYHAEEVQCDGR | AF-Q16527 |
| sp\|P23528\|COF1_HUMAN | AVLFCLSEDKK | 4BEX |
| sp\|P54105\|ICLN_HUMAN | LSWLDGSGLGFSLEYPTISLHALSR | AF-P54105 |
| sp\|P04792\|HSPB1_HUMAN | TKDGVVEITGK | 6DV5 |
| sp\|P14314\|GLU2B_HUMAN | SLKDMEESIR | AF-P14314 |
| sp\|Q14204\|DYHC1_HUMAN | KLVPLLLEDGGEAPAALEAALEEK | 5OWO |
| sp\|Q14204\|DYHC1_HUMAN | EWTDGLFTHVLR | 5NUG |
| sp\|O43143\|DHX15_HUMAN | YMTDGMLLR | 5XDR |
| sp\|P53621\|COPA_HUMAN | YAVTTGDHGIIR | AF-P53621 |
| sp\|Q8TEA8\|DTD1_HUMAN | SASSGAEGDVSSEREP | 2OKV |
| sp\|P52272\|HNRPM_HUMAN | MGLVMDR | AF-P52272 |
| sp\|P52272\|HNRPM_HUMAN | MAAPIDR | AF-P52272 |
| sp\|P62280\|RS11_HUMAN | EAIEGTYIDKK | 4UG0 |
| sp\|P62318\|SMD3_HUMAN | FLILPDMLK | 4PJ0 |
| sp\|P29401\|TKT_HUMAN | LILDSAR | 3MOS |
| sp\|Q12769\|NU160_HUMAN | SEDGEIVSTPR | 7R5K |
| sp\|P32119\|PRDX2_HUMAN | IGKPAPDFK | 1QMV |
| sp\|P55209\|NP1L1_HUMAN | YAVLYQPLFDKR | 7UN3 |
| sp\|P40926\|MDHM_HUMAN | MISDAIPELK | 2DFD |
| sp\|Q16891\|MIC60_HUMAN | ELDSITPEVLPGWK | AF-Q16891 |
| sp\|Q14566\|MCM6_HUMAN | VSGVDGYETEGIR | AF-Q14566 |
| sp\|P52701\|MSH6_HUMAN | LSDGIGVMLPQVLK | 2O8B |
| sp\|P42704\|LPPRC_HUMAN | TVLDQQQTPSR | AF-P42704 |
| sp\|P0DPH8\|TBA3D_HUMAN | LSVDYGK | AF-P0DPH8 |

Table S2. Isomerized peptides impacted by aging in Native lysis buffer compared to aging in RIPA lysis buffer. Crowding in both monomer and complexes are also shown.

| Protein | Peptide | Trend? | Monomer crowding | Complex crowding |
|---|---|---|---|---|
| sp\|P62826\|RAN_HUMAN | LVLVGDGGTGK | Increase | 9 | 9 |
| sp\|P62913\|RL11_HUMAN | YDGIILPGK | Decrease | 9 | 9 |
| sp\|P22234\|PUR6_HUMAN | IKAEYEGDGIPTVFVAVAGR | Decrease | 12 | 12 |
| sp\|P23284\|PPIB_HUMAN | TAWLDGK | Decrease | 9 | 9 |
| sp\|P50914\|RL14_HUMAN | ALVDGPCTQVR | Increase | 15 | 15 |
| sp\|P25398\|RS12_HUMAN | TALIHDGLAR | Decrease | 9 | 9 |
| sp\|P23396\|RS3_HUMAN | FVADGIFK | Increase | 14 | 14 |
| sp\|P23396\|RS3_HUMAN | ELAEDGYSGVEVR | Decrease | 10 | 10 |
| sp\|P52597\|HNRPF_HUMAN | VHIEIGPDGR | Decrease | 13 | 13 |
| sp\|P08238\|HS90B_HUMAN | YHTSQSGDEMTSLSEYVSR | Decrease | 13 | 13 |
| sp\|P32119\|PRDX2_HUMAN | ATAVVDGAFK | Decrease | 9 | 9 |
| sp\|P38646\|GRP75_HUMAN | SQVFSTAADGQTQVEIK | Decrease | 9 | 9 |
| sp\|Q06830\|PRDX1_HUMAN | ATAVMPDGQFK | Decrease | 12 | 12 |
| sp\|P55072\|TERA_HUMAN | IVSQLLTLMDGLK | Decrease | 14 | 14 |

Table S3. Isomerized peptides impacted by the addition of ammonium hydroxide to *in vitro* aging lysis buffers.

| Protein | Peptide | Structure | Trend | Condition |
|---|---|---|---|---|
| sp\|P23526\|SAHH_HUMAN | ESLIDGIKR | Alpha helix | Decrease | Both |
| sp\|P17066\|HSP76_HUMAN | TTPSYVAFTDTER | Flexible | Increase | Both |
| sp\|P26583\|HMGB2_HUMAN | SEHPGLSIGDTAK | Alpha helix | Increase | Both |
| sp\|P62937\|PPIA_HUMAN | KITIADCGQLE | Beta sheet | Decrease | Both |
| sp\|P32119\|PRDX2_HUMAN | ATAVVDGAFK | Flexible | Increase | Native |
| sp\|P55072\|TERA_HUMAN | IVSQLLTLMDGLK | Alpha helix | Increase | Native |
| sp\|P25705\|ATPA_HUMAN | VLSIGDGIAR | Flexible | Increase | Native |
| sp\|P08238\|HS90B_HUMAN | YHTSQSGDEMTSLSEYVSR | Flexible | Increase | Native |
| sp\|Q9Y536\|PAL4A_HUMAN | IIPGFMCQGGDFTR | Flexible | Decrease | RIPA |
| sp\|F5H284\|PAL4D_HUMAN | TEWLDGK | Flexible | Increase | RIPA |
| sp\|P61978\|HNRPK_HUMAN | VVLIGGKPDR | Alpha helix | Decrease | RIPA |
